# Intestinal Bacteria Maintain Adult Enteric Nervous System and Nitrergic Neurons via Toll-like Receptor 2-induced Neurogenesis in Mice

**DOI:** 10.1101/2020.03.01.968727

**Authors:** Shadi S. Yarandi, Subhash Kulkarni, Monalee Saha, Kristyn E. Sylvia, Cynthia L. Sears, Pankaj J. Pasricha

**Author notes:** **Correspondence** Pankaj Jay Pasricha, 720 Rutland Ave, Ross 958, Baltimore, MD. **Author contributions** S.S.Y, C.S. and P.J.P. conceived the study; S.S.Y, S.K., C.S., and P.J.P designed the research studies; S.S.Y and M.S. conducted the experiments; S.S.Y, J.G, K.S acquired and analyzed the data; S.S.Y, S.K. and P.J.P drafted the manuscript with input from all authors.

## Abstract

**Background & Aims:** The enteric nervous system (ENS) exists in close proximity to luminal bacteria. Intestinal microbes regulate ENS development, but little is known about their effects on adult enteric neurons. We investigated whether intestinal bacteria or their products affect the adult ENS via toll like receptors (TLRs) in mice.

**Methods:** We performed studies with conventional C57/BL6, germ-free C57/BL6, *Nestin*-creER^T2^:tdTomato, *Nestin*-GFP, and ChAT-cre:tdTomato. Mice were given drinking water with ampicillin or without (controls). Germ-free mice were given drinking water with TLR2 agonist or without (controls). Some mice were given a blocking antibody against TLR2 or a TLR4 inhibitor. We performed whole-gut transit, bead latency, and geometric center studies. Feces were collected and analyzed by 16S rRNA gene sequencing. Longitudinal muscle myenteric plexus (LMMP) tissues were collected, analyzed by immunohistochemistry, and levels of nitric oxide were measured. Cells were isolated from colonic LMMP of *Nestin*-creER^T2^:tdTomato mice and incubated with agonists of TLR2 (receptor for Gram-positive bacteria), TLR4 (receptor for Gram-negative bacteria), or distilled water (control) andd analyzed by flow cytometry.

**Results:** Stool from mice given ampicillin had altered composition of gut microbiota with reduced abundance of Gram-positive bacteria and increased abundance of Gram-negative bacteria, compared with mice given only water. Mice given ampicillin had reduced colon motility compared with mice given only water, and their colonic LMMP had reduced numbers of nitrergic neurons, reduced nNOS production, and reduced colonic neurogenesis. Numbers of colonic myenteric neurons increased after mice were switched from ampicillin to plain water, with increased markers of neurogenesis. Nestin-positive ENPCs expressed TLR2 and TLR4. In cells isolated from the colonic LMMP, incubation with the TLR2 agonist increased the percentage of neurons originating from ENPCs to approximately 10%, compared to approximately 0.01% in cells incubated with the TLR4 agonist or distilled water. Mice given an antibody against TLR2 had prolonged whole-gut transit times; their colonic LMMP had reduced total neurons and a smaller proportion of nitrergic neurons per ganglion, and reduced markers of neurogenesis compared with mice given saline. Colonic LMMP of mice given the TLR4 inhibitor did not have reduced markers of neurogenesis. Colonic LMMP of germ-free mice given TLR2 agonist had increased neuronal numbers compared with control germ-free mice.

**Conclusions:** In the adult mouse colon, TLR2 promotes colonic neurogenesis, regulated by intestinal bacteria. Our findings indicate that colonic microbiota help maintain the adult ENS via a specific signaling pathway. Pharmacologic and probiotic approaches directed towards specific TLR2 signaling processes might be developed for treatment of colonic motility disorders related to use of antibiotics or other factors.

## Introduction

The enteric nervous system (ENS), an autonomous nervous system contained entirely in the gastrointestinal tract wall, is critical for the maintenance of normal gut functions, including gut motility and secretion^1^. Developmental absence of enteric neurons, as seen in Hirschsprung’s disease, leads to significant disruption of colonic motility^2^, but much less is known about the pathogenesis of acquired motility disorders^3^. It has been assumed that such disorders may arise from injury or death of mature enteric neurons in what has traditionally been thought to be a static system with little or no capacity for renewal^4^. The ENS is continually exposed to a variety of extrinsic factors such as diet, mechanical stretch, medications, and toxins with recent data suggesting that many of these can influence the structure and function of ENS^5–9^. The mechanism through which these extrinsic factors can affect ENS are not fully understood, but the gut microbiota is emerging as one of the main mediators of these effects. Diet^10, 11^ or antibiotics^12–14^ can change the composition of the gut microbiota, which in turn may modulate the ENS^6,14, 15^

Our knowledge about the effects of gut microbiota on the ENS has largely been derived from studies in early development^14^. In germ-free (GF) mice, the ENS is both structurally and functionally abnormal^16, 17^ but improves with restoration of gut microbiota^17, 18^. These effects may be mediated by toll-like receptors (TLRs), which respond to bacterial cell wall products such as lipoteichoic acid (LTA) and lipopolysaccharide (LPS)^19^. Mice lacking TLR2, which preferentially binds to LTA, show a reduced number of ileal enteric neurons and a TLR2 agonist can ameliorate enteric neuronal loss when exposed to a cocktail of broad-spectrum antibiotics^20^. Similarly, mice deficient in TLR4 (the receptor for LPS), have abnormal development of ENS with reduced number of nitrergic neurons in colonic myenteric plexus^15^. Less is known about the effects of gut microbiota on the adult ENS. In adult mice with a fully developed ENS, reduction of gut microbes with a cocktail of antibiotics results in increased apoptosis and loss of enteric neurons ^15^, but the precise cellular target or the putative molecular mediators underlying this effect are not known.

We have recently shown that adult mature neurons in the small intestine degenerate and die, and are replaced by new neurons derived from Nestin-expressing enteric neuronal precursor cells (ENPCs)^21^. We hypothesized that bacterial products are important in maintaining the adult ENS via TLR-mediated effects on ENPCs. Here, we show that neurogenesis occurs in the adult colonic ENS. Inhibition of TLR2 signaling suppresses neurogenesis whereas activation of TLR2 promotes colonic neurogenesis from ENPCs in both antibiotic induced dysbiosis and germ-free murine models. Our results provide a mechanistic link between the microbiota and maintenance of a healthy ENS in adults and offer new insight into how dysbiosis can lead to disruption and dysfunction of this system.

## Materials and Methods

### Animals

All mice used for the experiments were between the ages of 8 and 16 weeks and are designated as adult mice. Experimental protocols were approved by the Johns Hopkins University’s Animal Care and Use Committee in accordance with the guidelines provided by the National Institutes of Health. Details of WT and transgenic mice used for the experiments are presented in SI Methods.

### Antibiotic treatment

8-16 weeks old C57/BL6 male mice were treated either with Ampicillin (0.5 g/L) through drinking water containing 4 g/L Splenda (Heartland food products group, Carmel, IN); or with Drinking Water containing 4 g/L Splenda without any antibiotics as a control group.

### Tamoxifen induction

Tamoxifen (Sigma-Aldrich) was dissolved in corn oil at a concentration of 20 mg/mL by shaking overnight at room temperature in a foil wrapped tube. For labeling the Nestin^+^ cells with tdTomato, 3 oral gavages of tamoxifen (100μL) was performed on consecutive days.

### Measuring GI motility

#### Measuring whole gut transit time

Whole gut transit time (WGTT) was measured at 1, 2, 3, and 4 weeks after the start of antibiotic treatment and 10 days after antibiotic discontinuation. Mice received 0.3 mL 6% carmine solution in 0.5% methylcellulose by oral gavage into the animal’s stomach as previously described ^21^. The time taken for each mouse to produce a red fecal pellet after the administration of carmine dye was recorded in minutes.

#### Determining the geometric center

Geometric center studies were performed 2 weeks after treatment. Distribution of the non-absorbable marker fluorescein isothiocyanate-dextran (FITC-dextran, Sigma) was determined in the colon of mice as described previously^22^.

#### Measuring distal colonic transit

Distal colonic transit time was measured using the bead latency test (details in SI methods).

### Bacterial DNA Extraction

Frozen fecal pellets were extracted using the DNeasy PowerSoil HTP 96 Kit (Qiagen, Venlo, Netherlands). One or two frozen fecal pellets were deposited in each well of the Powerbead plate of the Powersoil kit and extracted according to manufacturer’s instructions. Extracted DNA was quantified using the Quant-iT dsDNA high sensitivity kit (ThemoFisher, Waltham, MA).

### 16S rRNA gene PCR amplicon sequencing and library preparation

These methods are expanded in SI Methods.

### Tissue preparation and cell isolation

Tissue preparation and cell isolation protocols are described in SI Methods.

### Immunohistochemistry

Longitudinal muscle-containing myenteric plexus (LM-MP) was fixed in freshly made ice-cold 4% paraformaldehyde solution. The fixed myenteric plexus tissue was then washed twice in ice-cold PBS in the dark at 16 °C and then incubated in blocking-permeabilizing buffer (BPB; 5% normal goat serum with 0.3% Triton-X) for 1 h. After removing the tissue from the BPB, it was incubated with the primary antibody for HuC/D and nNOS for 48 h at 16 °C in the dark with shaking at 60 rpm. The tissue was then washed three times in PBS at room temperature in the dark and incubated in the appropriate secondary antibody at room temperature for 1 h while on a rotary shaker (65 rpm). The tissue was again washed three times in PBS at room temperature, counterstained with DAPI to stain the nuclei, overlaid with Vectashield mounting medium and cover-slipped. Slides were imaged using a Leica 510 confocal microscope and analyzed using Fiji software. The number of neurons per ganglion^21, 23, 24^ was enumerated to adjust for changes in the size of cecum in antibiotic treated mice and GF mice and ANOVA was used to compare the number of neurons per ganglion between the groups. Details of reversibility, neurogenesis experiments and experiments in GF mice are explained in SI methods.

### Antibodies, drugs and concentrations

All of the antibodies sources and concentrations are listed in table S1. Agonists, antagonists and inhibitors sources and concentrations are listed in table S20.

### Measurement of nNOS mediated NO release

LM-MP from ampicillin, control and withdrawal groups were washed in standard Krebs solution and then incubated for 1 hour at 37 °C with or without Nomega-Nitro-L-arginine (L-NNA), a specific inhibitor for nNOS^25–27^. At the end of the incubation period, the supernatants were collected and used for NO quantification. Nitric oxide (NO) levels were determined by the modified Griess method using a colorimetric kit (Abcam, Cambridge, MA), according to the manufacturer’s instructions. Absorbance of samples was determined at 540 nm in an automated microplate reader. Results were compared with a standard curve generated with sodium nitrate. The nNOS-generated fraction of the total NO released was calculated by subtracting the amount of nitrate in the LNNA group (non-nNOS generated) from the total nitrate using the formula, 100− (100 × L-NNA/no L-NNA). Data were expressed as mean ± SEM.

### *In vitro* differentiation assay

Nestin-creER^T2^:tdTomato mice post-tamoxifen induction were sacrificed and myenteric plexus were isolated and dissociated in a digestion buffer as described above. Cells obtained from myenteric plexus were then plated and cultured for 14 days at the presence of TLR2 agonist; Pam3Csk, TLR4 agonist; LPS, or placebo. After 14 days, cells were fixed in 2% PFA for 15 minutes, washed, blocked with 2% BSA (Sigma), treated with digestion buffer as above for 20 min at 22°C and then stained with antibodies against HuC/D and nNOS (Table S1) diluted in permeabilization buffer. Cells were then studied by Flow analyses to study the expression of HuC/D and nNOS.

### Flow analyses

Flow analyses to study the expression of TLR2, HuC/D, and nNOS in Nestin derived neurons was performed on BD LSR II.UV (See SI methods for more details). All analyses of FACS data were performed using FlowJo 7.6.1.

### Statistics

All the quantified data is presented in Tables S2–S19. Data are expressed as the mean ± Standard Error for every graphical representation. Statistical analysis was performed with SPSS (SPSS 23, IBM, 2015). ANOVA test were used to compare means for statistically significant difference between groups. Statistical significance (*) was assumed if p < 0.05. Details of microbiome statistical analyses are explained in SI Methods.

### Data Generation

Sample sizes of numbers of animals and ganglia counted are shown in tables S2–S19. In all experiments, littermates were used, and in cases where more numbers of mice were required, we used mice of the same age that were of the same genotype. Mice from different litters were then distributed equally between different groups. For microscopic counts, random fields were imaged and the numbers of cells per ganglia were enumerated per group. For FACS analysis, 10,000 cells were assayed. All *in vitro* experiments were performed three times. All *in vivo* experiments were performed on 3-5 mice per group, and data were generated using technical replicates of every mouse, as denoted above.

## Results

### Neurogenesis in the adult colonic myenteric plexus is driven by Nestin^+^-neuronal precursors

We have previously identified Nestin as a marker of the ENPCs in the small intestine and showed that a subset of Nestin^+^ cells give rise to neurons *in vivo*^21^. Nestin^+^ cells also exist in the colon and surround the neurons within the colonic myenteric ganglion (Figure S1A). In inducible Cre transgenic mice (Nestin-creER^T2^:tdTomato) the pan-neuronal marker HuC/D did not show overlap between tdTomato-expressing cells and neurons at 12 hours after tamoxifen induction(Figure S1B(i)); but after a subpopulation of tdTomato-expressing neurons emerged within the colonic myenteric plexus, indicating their derivation from Nestin-expressing cells (Figure S1B(ii)).

### ENPCs express TLR2 and TLR4

Both TLR2 and TLR4 were highly expressed by Nestin-GFP^+^ cells isolated from the colonic myenteric plexus of reporter mice (97.7% and 88.2%, respectively). We then examined temporal changes in the expression of these receptors by giving tamoxifen to Nestin-creER^T2^:tdTomato mice 3 days before sacrifice, after which cells from the colonic myenteric plexus were isolated and cultured for 14 days. The expression of TLR2 and TLR4 remained high in newly born neurons identified as HuC/D^+^ tdTomato^+^ cells (75% and 89%, respectively), but much lower in older neurons, identified as HuC/D^+^ tdTomato^−^ cells (21.59% and 47%, respectively) (Figure 1A-B).

**Figure 1.**
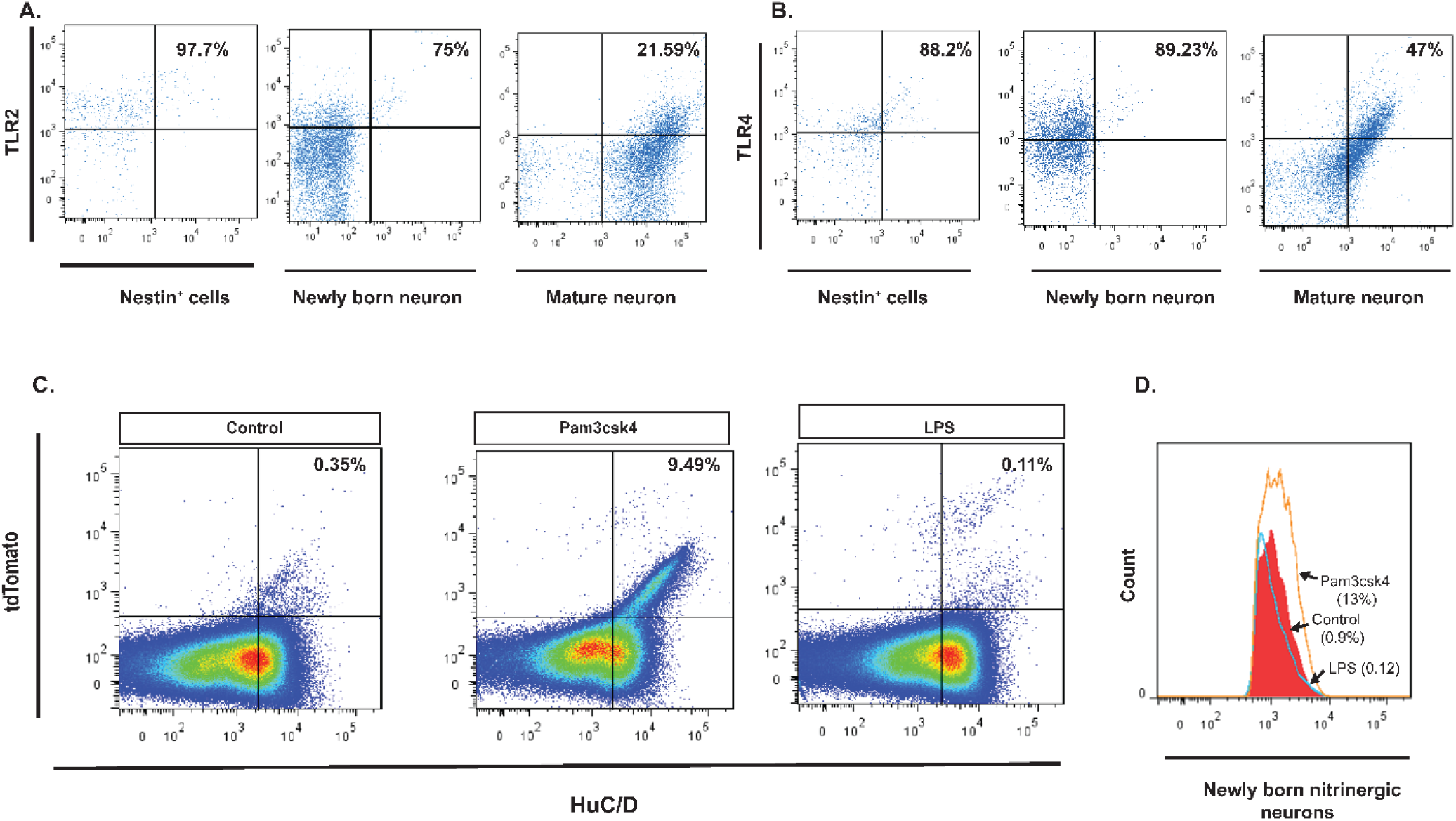
Both TLR2 and TLR4 are expressed on Nestin^+^ ENPCs and existing neurons but only TLR2 signaling increases Nestin^+^ ENPC driven neurogenesis in vitro. (A. left panel) Flow cytometry of Nestin^+^ ENPCs isolated from colonic myenteric plexus of Nestin-GFP mice and stained with TLR2, showing that 97.7% of Nestin^+^ cells express TLR2 (top right quadrant). (A. middle panel) Flow cytometry of Huc/D^+^ isolated from colonic myenteric plexus of Nestin-creER^T2^:tdTomato mice after tamoxifen induction and stained with TLR2, showing that 75% of HuC/D^+^tdTomato^+^ neurons express TLR2. (A. right panel) Flow cytometry of HuC/D^+^ neurons that do not express tdTomato shows that 21.59% of these cells express TLR2. (B. left panel) Flow cytometry of Nestin^+^ ENPCs isolated from colonic myenteric plexus of Nestin-GFP mice and stained with TLR4, showing that 88.2% of Nestin^+^ cells express TLR4. (B. middle panel) Flow cytometry of Huc/D^+^ isolated from colonic myenteric plexus of Nestin-creER^T2^:tdTomato mice after tamoxifen induction and stained with TLR4, showing that 89.2% of HuC/D^+^tdTomato^+^ neurons express TLR4. (B. right panel) Flow cytometry of HuC/D^+^ neurons that do not express tdTomato shows that 47% of these cells express TLR4. (C) Pseudocolor plots of flow cytometry data of cells isolated from colonic myenteric plexus of Nestin-creERT2:tdTomato mice after tamoxifen induction and being cultured for 14 days and stained for HuC/D. (C. left panel) Flow cytometry from control culture group showing newly formed neurons from Nestin^+^ ENPCs that express tdTomato (top right quadrant). (C. middle panel) Addition of TLR2 agonist (Pam3csk4) increased the number of newly formed neurons. (C. right panel) Addition of TLR4 agonist (LPS) did not increase the population of newly formed neurons. (D) Flow cytometry of HuC/D^+^nNOS^+^ cells isolated from colonic myenteric plexus of Nestin-creER^T2^:tdTomato mice and cultured for 14 days with addition of Pam3csk4 (orange line), LPS (blue line), or vehicle (red filled curve) showing increase in the population of newly formed nitrinergic neurons with TLR2 agonist addition.

### Activation of the TLR2 signaling pathway increases neurogenesis *in vitro*

We next cultured the cells isolated from the colonic myenteric plexus of Nestin-creER^T2^:tdTomato for 14 days in the presence of Pam3CSK4 (a synthetic triacylated lipopeptide that activates the TLR2/TLR1 heterodimer), LPS (lipopolysaccharide, a natural ligand for TLR4) or vehicle. FACS analysis showed that the percentage of HuC/D^+^tdTomato^+^ neurons (indicating an origin from Nestin^+^ precursors) was increased from ~0.01% in the control group to ~10% in cells treated with Pam3CSK4 (Figure 1C) while it was unchanged in the LPS group. Further, the proportion of tdTomato^+^ neurons expressing nNOS increased from ~0.9% in control to ~13% in cells treated with Pam3CSK4, but remained the same in the LPS group (Figure 1D). These data indicate that activation of TLR2, but not TLR4, promotes the development of neurons that are enriched in nNOS.

### Inhibition of the TLR2 signaling decreases neurogenesis from ENPCs *in vivo*

T2.5 (a blocking anti-TLR2 monoclonal antibody ^28^) was administered to Nestin-creER^T2^:tdTomato mice, along with tamoxifen. Whole gut transit was significantly prolonged in mice treated with T2.5 as compared with controls, accompanied by a significant decrease in the total number of HuC/D^+^ neurons as well as the proportion of nitrergic neurons per ganglion (Figure 2). The proportion of neurons expressing tdTomato per ganglion (number of HuC/D^+^ tdTomato^+^ neurons/total number of HuC/D^+^ neurons) was also significantly lower in treated mice (Figure 2A-B and D), indicating suppression of neurogenesis.

**Figure 2.**
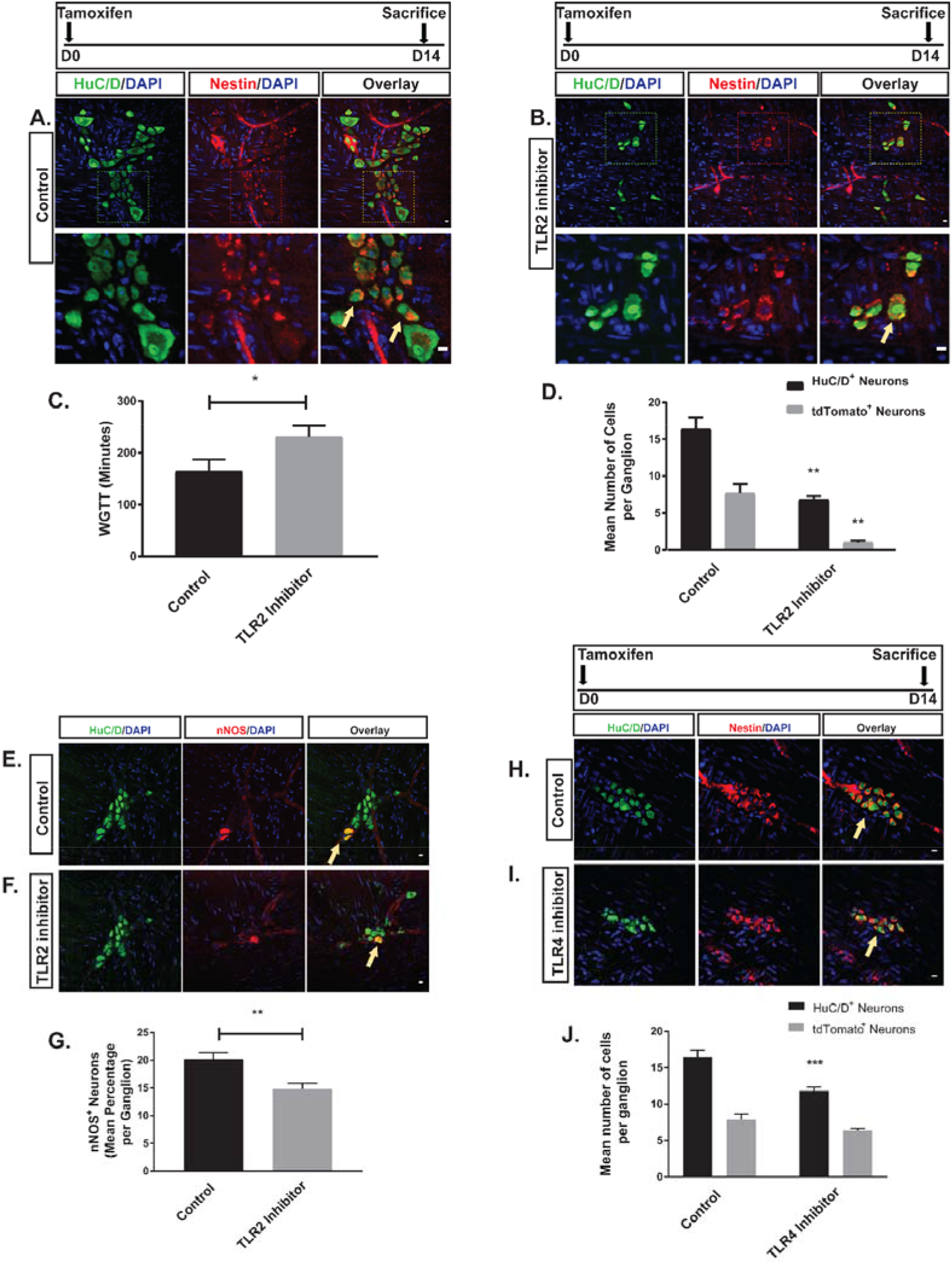
Effects of TLR2 and TLR4 signaling on colonic neurogenesis. A-B are representative photomicrographs of colonic myenteric ganglion of Nestin-creER^T2^:tdTomato mice after tamoxifen induction and immunostaining with the neuronal marker HuC/D (green) in two groups of vehicle and TLR2 inhibitor treated animals. E panels are representative photomicrographs of colonic myenteric ganglion after immunostaining with the neuronal marker HuC/D (green) and nNOS (red) in the same groups. (A) tdTomato^+^ neurons (arrow) within the myenteric plexus ganglion of control mice. Top panel and bottom panel overlays represent different magnifications showing that nestin derived neurons in the myenteric plexus (arrow) is reduced in mice treated with T2.5 (B). (C) Grouped results of WGTT under various conditions shows that WGTT is prolonged in the TLR2 inhibitor group (*p<0.05). (D) Grouped results of mean number of neurons and tdTomato–expressing neurons (newly formed neurons from ENPCs) under different conditions (**p<0.01). (E) Representative image of normal colonic myenteric ganglion containing nitrergic neurons (arrow). (F) T2.5 treatment reduced the total number of enteric neurons in a ganglion and reduced the proportion of nitrergic neurons within the ganglion. (G) Grouped results of mean percentage of nitrergic neurons under various conditions (**p<0.01). H-I are representative photomicrographs of colonic myenteric ganglion of Nestin-creER^T2^:tdTomato mice after tamoxifen induction and immunostaining with the neuronal marker HuC/D (green) in two groups of vehicle and TLR4 inhibitor treated animals. tdTomato^+^ neurons (arrow) within the myenteric plexus ganglion of control mice (H) and TLR4 inhibitor treated mice (I). (J) Grouped results of mean number of neurons and tdTomato–expressing neurons (newly formed neurons from ENPCs) under different conditions (***p<0.001). Scale bar is 10 μm.

In order to identify the role of TLR4 in modulating neurogenesis, the TLR4 antagonist C34 ^29^ was administered to the Nestin-creER^T2^:tdTomato mice via the oral route at a concentration of 1 mg/kg at the same time as tamoxifen induction and continued for 14 days. After 14 days, cecal LMMP preparations were stained for HuC/D. There was a decrease in the total number of neurons per ganglion after C34 treatment, but this effect was not accompanied by a significant decrease in the proportion of neurons expressing tdTomato (Figure 2H-J).

### Suppression of gram-positive bacteria in the gut decreases neurogenesis and results in alteration of ENS structure and function in a reversible manner

Since our *in vitro* experiments did not indicate an effect of LPS we hypothesized that gram-positive bacteria may play a more important role in colonic neurogenesis. We therefore investigated the effects of oral administration of ampicillin, a beta-lactam antibiotic that preferentially targets gram-positive bacteria.

Oral ampicillin administration (in drinking water mixed with Splenda) to conventional adult C57BL/6 mice significantly prolonged whole gut transit time (WGTT) as compared with controls (Figure 3), an effect that took 2 weeks to develop. In separate experiments, we showed that Splenda by itself does not affect WGTT (Figure S2). By contrast to the WGTT, bead expulsion latency time, a measure of distal colonic transit time (DCTT), was similar in ampicillin and control groups (Figure S3). Geometric center analysis of FITC-dextran dye transit assay showed that transit was delayed predominantly in the proximal colonic region of ampicillin-treated animals (Figure 3B). After ampicillin was discontinued, WGTT normalized by 10 days (Figure 3C).

**Figure 3.**
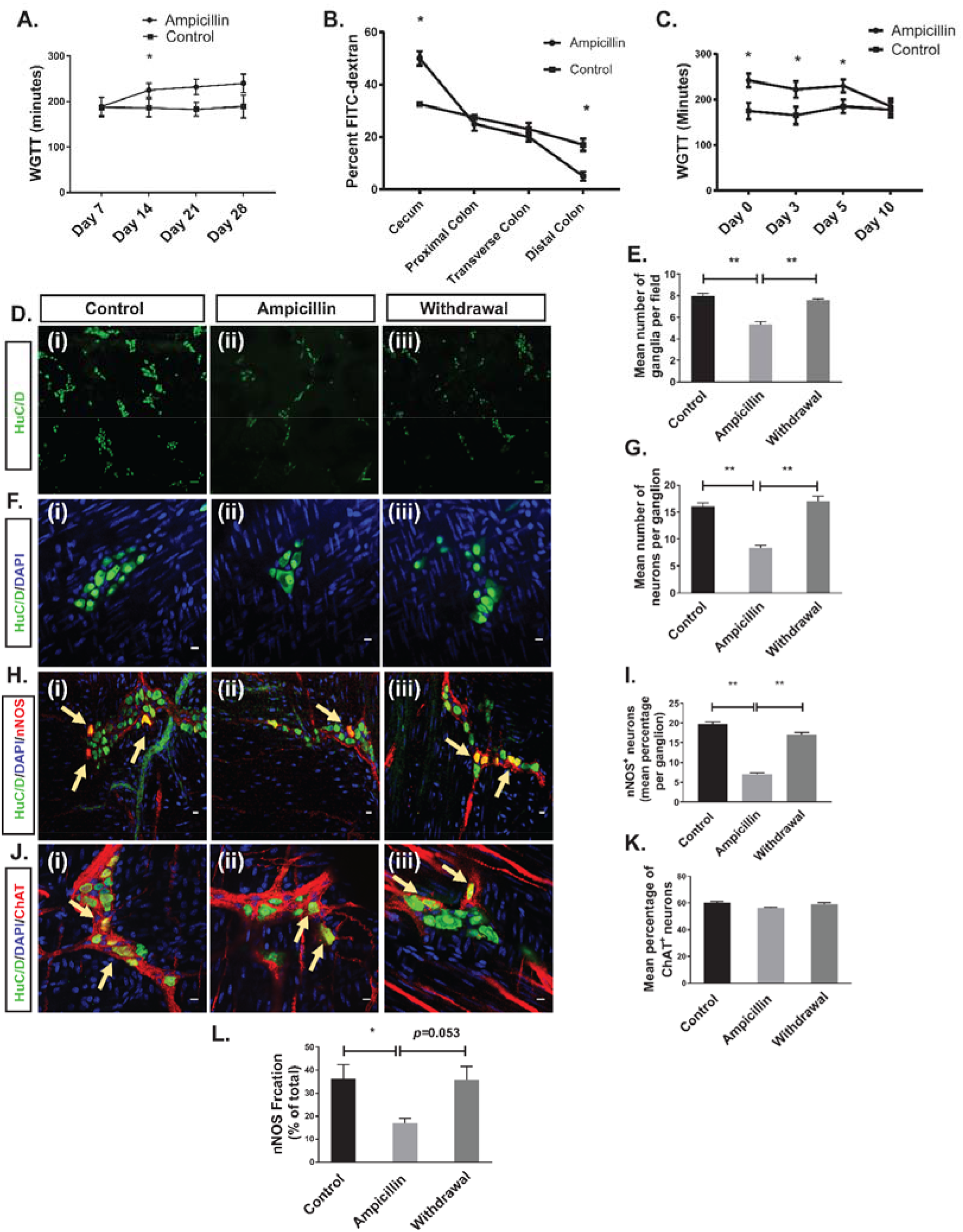
Ampicillin slows the proximal colon transit time reversibly through reduction of the ganglionic density, neuronal numbers/ganglia, and percentage of nitrergic neurons causing a decline in nitric oxide synthase (nNOS)-mediated NO release. (A) Ampicillin prolonged whole gut transit time after 2, 3, and 4 weeks of treatment. (B) FITC dextran test showed that the slowing of transit time occurs predominantly in the cecum. (C) WGTT normalized 10 days after discontinuation of ampicillin. (*p<0.05). D (i-iii) and F (i-iii) are representative photomicrographs of colonic myenteric plexus after immunostaining with the pan-neuronal marker HuC/D (green); nuclei are stained with DAPI (blue). H (i-iii) are representative photomicrographs of a colonic myenteric ganglion after immunostaining with the neuronal marker HuC/D (green) and nitrergic neuron marker nNOS (red); nuclei are stained with DAPI (blue). J (i-iii) are representative images of the colon wall from a ChAT-cre:tdTomato mouse where myenteric cholinergic neurons express red fluorescent protein tdTomato (red) after immunostaining with the neuronal marker HuC/D. Nuclei are counter stained with DAPI. (D (i)) Immunostaining with antisera against HuC/D (green) of myenteric plexus in control mice shows normal ganglion density. (D (ii)) Ampicillin treatment reduced the ganglion density. (D (iii)) Ganglion density recovered 10 days after discontinuation of ampicillin. (E) Grouped results of ganglion density under various conditions (**p<0.001). (F (i)) Immunostaining with antisera against HuC/D shows neurons in a representative colonic myenteric ganglion in control mice. (F (ii)) Ampicillin treatment reduced the number of neurons per ganglion. (F (iii)) 10 days after discontinuation of ampicillin, the number of neurons per ganglion normalized. (G) Grouped results of neuronal number per ganglion (** p<0.001). (H (i)) Representative image of a normal myenteric ganglion including nitrergic neurons (arrow). (H (ii)) Ampicillin treatment reduced the number and percentage of nitrergic neurons (H (iii)) Number of nitrergic neurons (arrows) and percentage of nitrergic neurons recovered 10 days after discontinuation of ampicillin. (I) Grouped results of percentage of nitrergic neurons per ganglion under various conditions (*p<0.05). (J (i)) Representative image of a normal myenteric ganglion including cholinergic neurons (arrow). (J (ii)) Ampicillin treatment did not reduce the percentage of cholinergic neurons (J (iii)) Percentage of cholinergic neurons did not change 10 days after discontinuation of ampicillin. (K) Grouped results of percentage of cholinergic neurons per ganglion under various conditions (p=0.31). (L) Mean nNOS percentage of the total NO released by the LM-MP isolated from the cecum in three different groups of control, ampicillin and withdrawal (*P < 0.01). Scale bar is 100 μm in D panel and 10 μm in F, H and J panel.

Ampicillin treatment resulted in a significant decrease in ganglionic density, the average number of neurons per ganglion and the proportion of nitrergic neurons (Figure 3D-I). However, the proportion of cholinergic neurons remained unchanged when ampicillin was given to ChAT-cre:tdTomato mice in which cholinergic neurons express red fluorescent protein tdTomato^21^ (Figure 3J-K). These changes were not accompanied by evidence of epithelial inflammation or injury (Figure S4). As with colonic transit, the neuronal changes returned to normal 10 days after discontinuing ampicillin (Figure 3D-I). We also measured the nNOS-generated fraction of total NO release from cecal LM-MP preparations. Ampicillin treatment resulted in decreased nNOS-mediated NO release, with a trend toward normalization 10 days after ampicillin discontinuation (Figure 3L).

Cecal microbiota of mice on ampicillin treatment for 14 days had significantly decreased diversity of microbial population as compared with the control (vehicle treated) group. Principal coordinate analysis (PCoA) showed reduced dimensionality of the microbial population in the ampicillin group (Figure 4A) with a significant reduction in Shannon alpha diversity index (Figure 3B). Both of these changes reversed 10 days after stopping ampicillin (Figure 4A-B).

**Figure 4.**
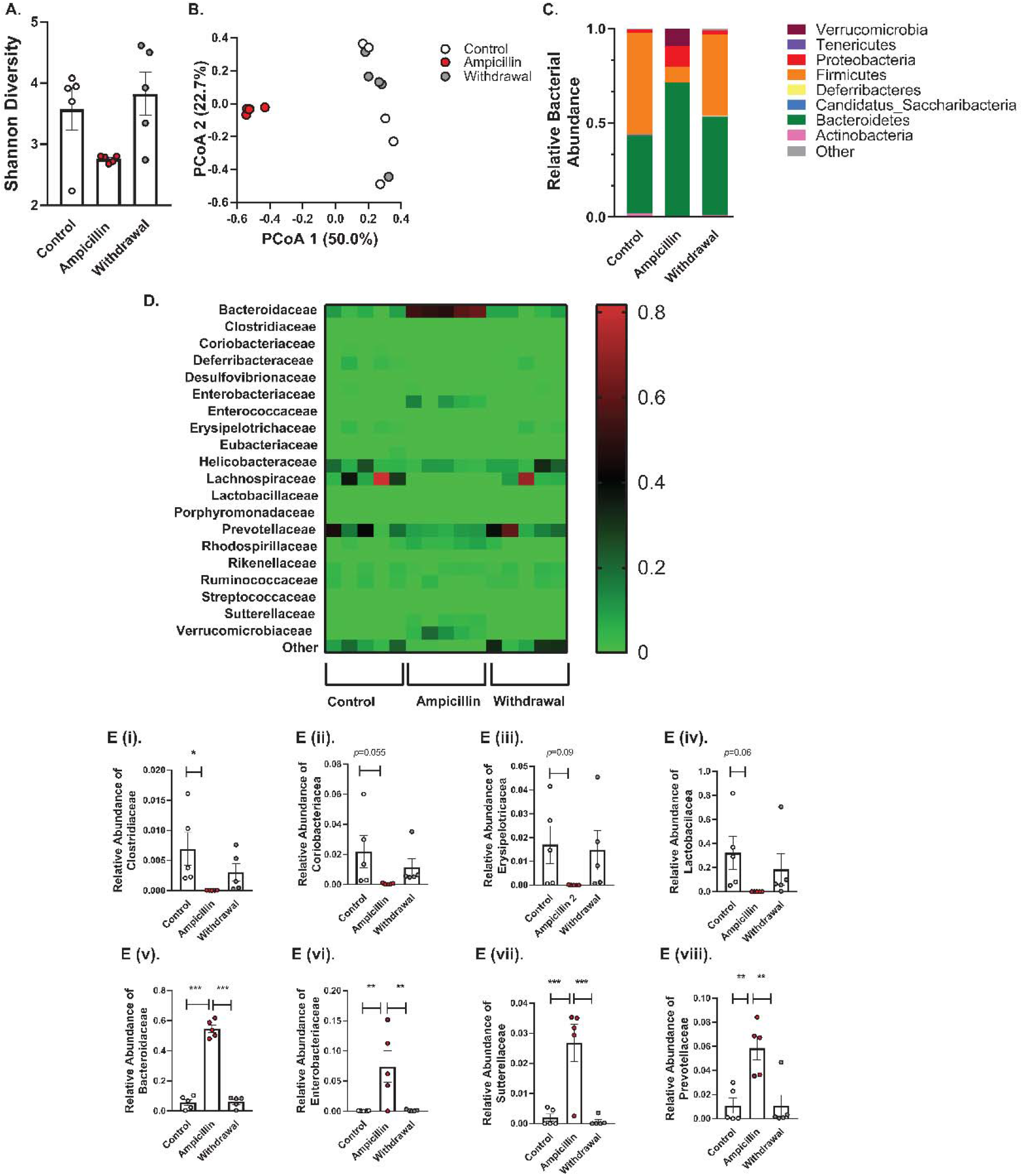
Ampicillin treatment creates a dysbiosis state in the cecum with a tendency toward normalization 10 days after ampicillin discontinuation. (A) Shannon diversity comparisons showed that alpha diversity decreased with ampicillin and increased 10 days after ampicillin discontinuation. (B) PCoA analysis using the weighted UniFrac suggested that the overall community composition from ampicillin group was different from control group or withdrawal group. (C) At the phylum level, treatment with ampicillin caused a significant increase in relative abundance of Bacteroidetes, Proteobacteria and Verrucomicrobia and a significant decrease in Firmicutes and Actinobacteria, these changes were reversible after 10 days of ampicillin discontinuation. (D) Heatmap of differentially abundant OTUs in different groups at family level. E. Relative bacterial abundance at family level. The relative abundance of E (i) *Clostridiaceae,* E (ii) *Coriobacteriacea*, E (iii) *Erysipelotricacea*, and E (iv) *Lactobacilacea* and had a trend toward increase after ampicillin discontinution. E (v) *Bacteroidaceae*, E (vi) *Enterobacteriaceae*, E (vii) *Suterellaceae*, and E (viii) showed significant increase in relative abundance after ampicillin treatment, an effect which was reversed 10 days after stopping ampicillin.

Ampicillin treatment resulted in an increase in the relative abundance of gram-negative microbes (*Bacteroidetes*, *Proteobacteria* and *Verrucomicrobia* at phylum level) and a decrease in that of gram-positive microbes (*Firmicutes* and *Actinobacteria at Phylum level* and *Clostridiaceae* at family level) (Figure 4C). Other families of gram-positive microbes also showed a trend towards a reduction with ampicillin treatment (Figure 4E (i-iv)). In contrast, 10 days after stopping ampicillin, the relative abundance of gram-positive families started increasing whereas that of gram-negative microbes declined 10 days after stopping the ampicillin (Figure 4E (v-viii)).

We next examined the effects of ampicillin-induced dysbiosis on neurogenesis in the cecum. Nestin-GFP^+^ reporter mice treated with ampicillin for two weeks showed a significant increase in the number of Nestin^+^ cells within myenteric ganglia (Figure 5A-C). In the presence of reduced numbers of mature neurons, this indicated that ampicillin treatment impaired neuronal differentiation rather than ENPC proliferation. This was further examined in Nestin creER^T2^:tdTomato mice that were treated with ampicillin for 14 days, with tamoxifen induction on day 8 (according to the experimental plan shown in Figure 5). There was a significant decrease in the number tdTomato^+^ neurons in ampicillin-treated mice as compared to controls (Figure 5D and 5F).

**Figure 5.**
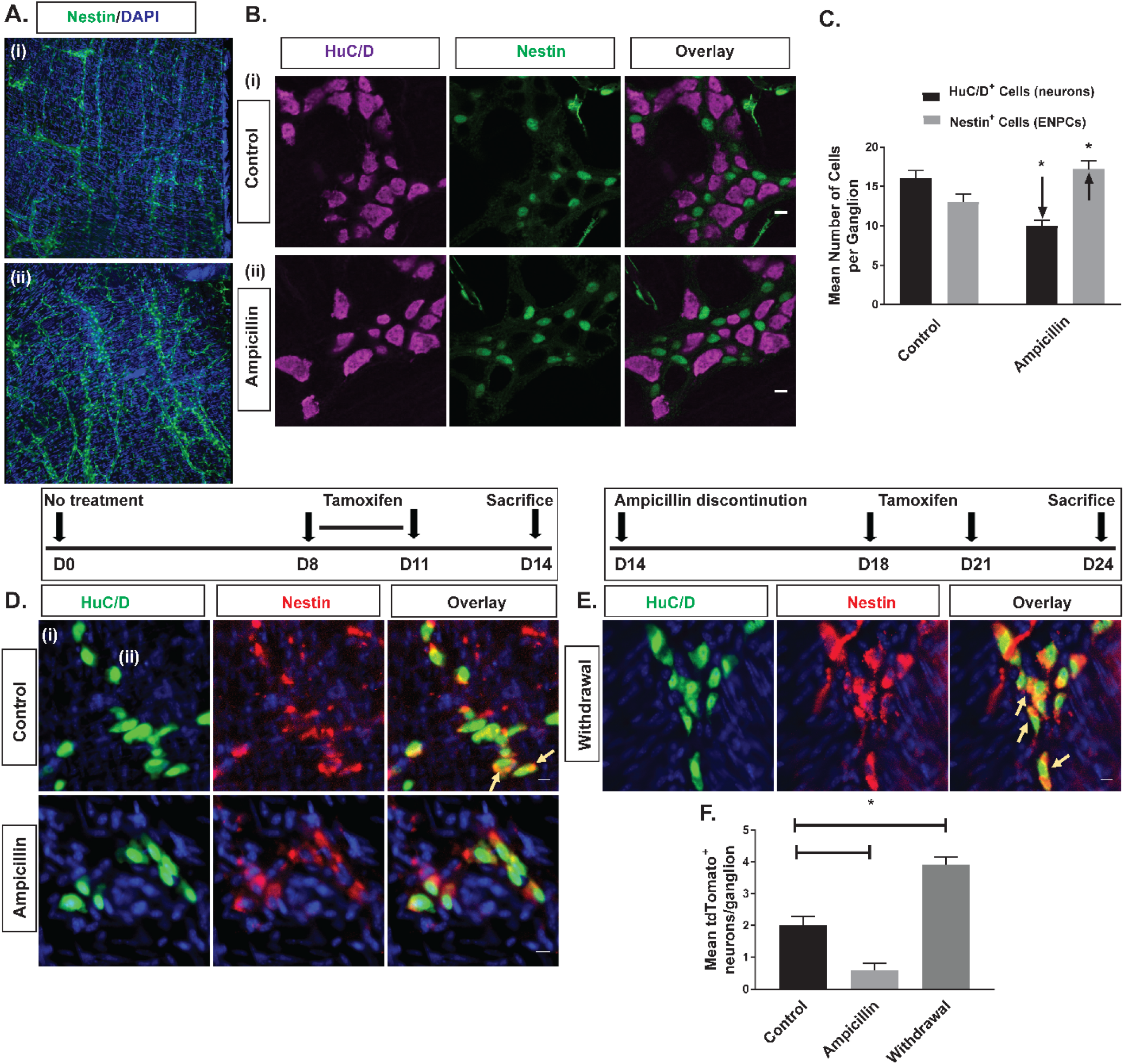
Neurogenesis from Nestin^+^ ENPCs was reduced by ampicillin despite increased number of Nestin-GFP^+^ cells within the colonic myenteric plexus. A (i-ii) are representative images of the colon wall from a Nestin-GFP reporter mouse using a 10x objective, nestin^+^ cells express GFP and the nuclei are counterstained with DAPI (blue); B (i-ii) are representative photomicrographs of Nestin-GFP mice after immunostaining with the neuronal marker HuC/D (purple). D-F are representative photomicrographs of colonic myenteric ganglion of Nestin-creER^T2^:tdTomato mice 6 days after tamoxifen induction and after immunostaining with the neuronal marker HuC/D (green); (A(i)) cells expressing Nestin-GFP (green) form an extensive network that is present in full thickenss colon wall. (A(ii)) Ampicillin treatment increases the number of Nestin-GFP cells in the myenteric plexus. (B(i)) within the colonic myenteric ganglion of Nestin-GFP mice, Nestin expressing cells (green) surround the HuC/D^+^ neurons. (B(ii)) Treatment with ampicillin results in increased number of cells expressing Nestin-GFP, while reduced the number of HuC/D^+^ neurons. (C) Grouped results of number of Nestin^+^ cells under various conditions. (* p<0.05). (D(i)) Newly formed neurons (arrow) are observed in the myenteric ganglion of control mice (D(ii)) Ampicillin treatment reduced the number of newly-formed neurons in the myenteric ganglion. (E) 10 days after discontinuing ampicillin, myenteric ganglia are filled with newly formed neurons (arrows). (F) Grouped results of mean number of newly formed neurons (expressing tdTomato) per ganglion under various conditions. (* p<0.05). Scale bar is 10 μm.

A separate cohort of mice were sacrificed 10 days after discontinuation of ampicillin (Withdrawal group) with tamoxifen induction initiated on day 6 prior to sacrificing this cohort. In the Withdrawal group, there was a significant increase in tdTomato+ neurons as compared with mice on ampicillin (Ampicillin group), and those who never received ampicillin (Control group) suggesting a reboundin neurogenesis (Figure 5D-F).

To ensure that ampicillin did not have a direct toxic effect on enteric neurons, we added ampicillin to cells isolated from the myenteric plexus of Nestin-creER^T2^:tdTomato mice and found that it did not reduce the number of neurons or number of tdTomato^+^ neurons *in vitro* (Figure S5.A). Additionally, we treated GF mice with ampicillin for 2 weeks and found no differences in gut motility or enteric neuronal number (Figure S5.B).

### TLR2 activation rescues ampicillin-induced changes in colonic ENS structure and function by increasing neurogenesis *in vivo*

Adult C57BL/6 mice were treated with the naturally occurring TLR2 agonist, LTA (Ampicillin-LTA) by daily oral gavage, or vehicle along with ampicillin (Ampicillin-vehicle) for 2 weeks; and compared them to a no-treatment (vehicle-only) control group. Whole gut transit in Ampicillin-LTA mice was not significantly different from the control group but was significantly faster as compared with the Ampicillin-vehicle group, indicating that concurrent treatment with LTA prevented the effects of ampicillin on WGTT (Figure 6D). Immunohistochemical studies of the colonic myenteric plexus of mice showed that HuC/D^+^ neuronal numbers and the proportion of nitrergic neurons were significantly higher in the Ampicillin-LTA group and similar to the no-treatment group (Figure 6A-C and 6E).

**Figure 6.**
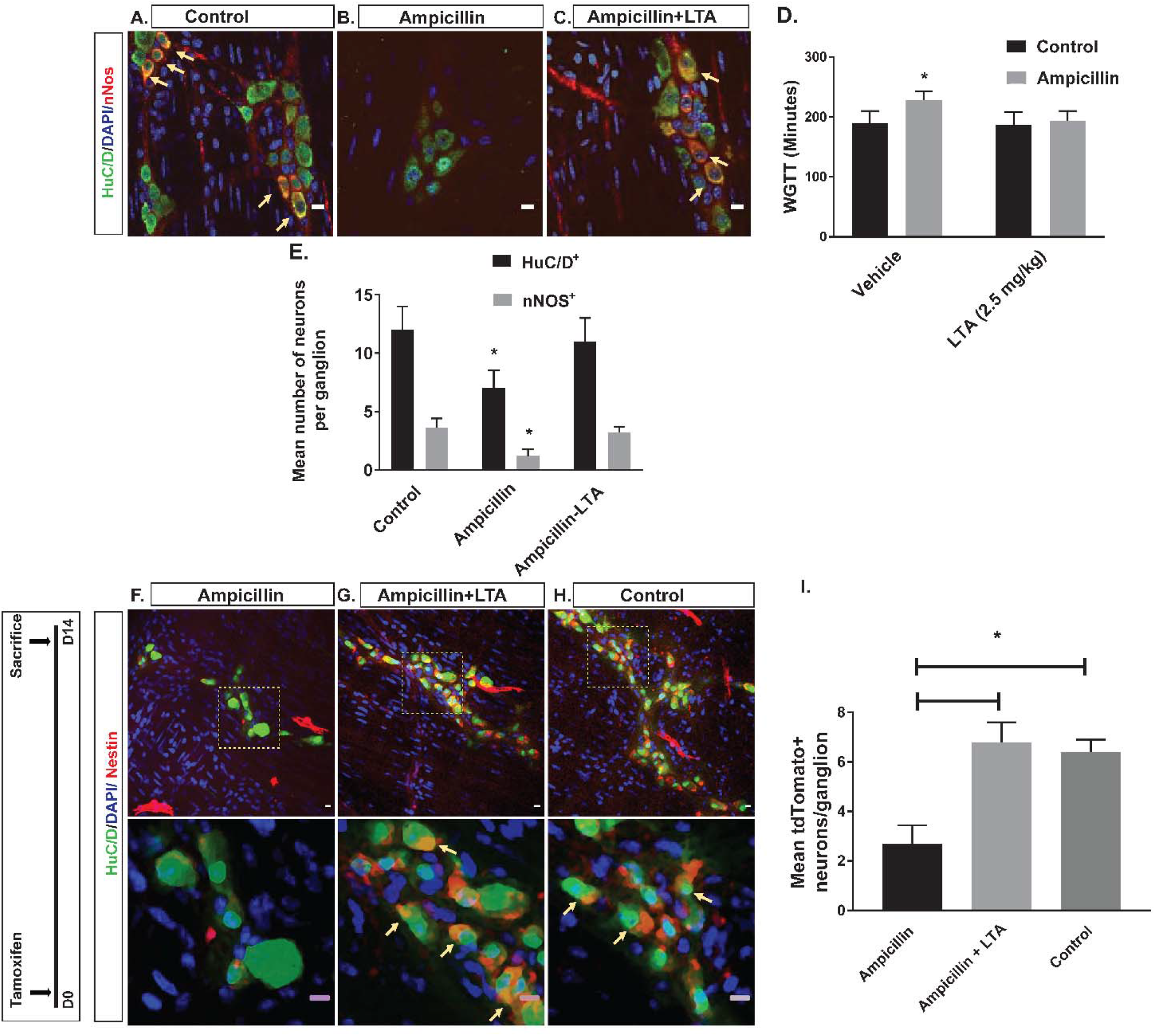
TLR2 signaling prevents ampicillin-induced dysmotility and loss of colonic neurons through enhancement of Nestin^+^ derived neurogenesis. A-C are representative photomicrographs of colonic myenteric ganglion after immunostaining with the neuronal marker HuC/D (green) and nNOS (red); nuclei are stained with DAPI (blue); (A) Representative image of normal colonic myenteric ganglion containing nitrergic neurons (arrow). Ampicillin reduced the total number of enteric neurons in a ganglion and reduced proportions of nitrergic neurons within the ganglion. (C) Concurrent treatment with LTA and ampicillin preserved the number of enteric neurons as well as proportions of nitrergic neurons (arrow) within myenteric ganglion. (D) Grouped results of WGTT under various conditions shows that WGTT is preserved in the ampicillin-LTA group. (E) Grouped results of mean number of total neurons and nitrergic neurons under various conditions shows preserved number of all neurons and nitrergic neurons in the ampicillin-LTA group. F-H are representative photomicrographs of colonic myenteric ganglion of Nestin-creER^T2^:tdTomato mice after tamoxifen induction and after immunostaining with the neuronal marker HuC/D (green). Top panels and bottom panels show different magnifications; (F) In ampicillin-treated mice, number of neurons expressing tdTomato (arrow) after tamoxifen induction is reduced, suggesting reduced neurogenesis. (G) Concurrent treatment with LTA and Ampicillin reverses this effect and increases the number of neurons expressing tdTomato (arrow), suggesting increased neurogenesis from Nestin^+^ ENPCs. (H) tdTomato^+^ neurons (arrow) within the myenteric plexus ganglion of control mice, showing neurogenesis from Nestin^+^ ENPCs at steady state. (I) Grouped results of mean number of tdTomato–expressing neurons (newly formed neurons from ENPCs) under different conditions (* p<0.05). Scale bar is 10 μm.

We next investigated the effects of TLR2 agonists on neurogenesis *in vivo* by quantifying the number of newborn neurons within the colonic myenteric plexus in Nestin-creER^T2^:tdTomato mice. The proportion of tdTomato^+^ neurons amongst all neurons per ganglion (number of HuC/D^+^ tdTomato^+^ neurons/total number of HuC/D^+^ neurons) was significantly higher in the Ampicillin-LTA group compared to the Ampicillin-vehicle group (Fig. 6F-I). The rate of neurogenesis in mice with concurrent LTA and ampicillin treatment was similar to that of the control group (Figure 6F-I), indicating that LTA enhances neurogenesis from Nestin^+^ cells.

### TLR2 activation also restores ENS structure in germ-free (GF) mice

We next used GF mice, a model of enteric neuropathy caused by a lack of microbiota. In agreement with previous reports on the small intestine of GF mice,^30^ we observed that untreated adult GF (GF-control) mice had reduced ganglionic density, as well as reduced numbers of myenteric neurons per ganglion in the colon (Figure S6A-D and S6G-H) and that this loss of neurons disproportionately affected nitrergic neurons (Figure S6E-F and Figure S6I).

We then treated GF mice with LTA or vehicle through drinking water for different durations of 2 or 3 weeks. Before sacrifice, stool was collected and cultured to ensure maintenance of the GF state. In GF mice treated with LTA for 2 weeks, the mean number of neurons per ganglion showed a significant increase compared to controls, but the percentage of nitrergic neurons was not statistically different (Figure S6L and Figure S6O). However, after 3 weeks of treatment with LTA, both of these measures were significantly higher compared to controls, indicating that TLR2 activation can promote neurogenesis in GF mice and push them towards a nitrergic fate (Figure S6J-O).

## Discussion

Our understanding of the maintenance of the adult ENS is limited and until recently constrained by the dogma that it is a static system with a stable population of neurons through most of life^4^. The recent demonstration that the ENS of the small intestine is in a state of constant turnover with active neurogenesis required to replace continually dying neurons^21^ has highlighted the vulnerability of this system and the importance of understanding the underlying mechanisms that maintain it. The gut microbiota, because of its abundance, proximity and known role in the development of the ENS^16, 17, 30^ is expected to be an important participant in this process. Microbial products can potentially access the intramural constituents of the colonic wall directly or exert their effects via indirect mechanisms involving epithelium, macrophages or other cell types with access to the lumen^15, 31, 32^ and we hypothesized that some of these products are important to maintain the colonic ENS, specifically via effects on neurogenesis.

In this study, we first showed that similar to their counterparts in the small intestine, the myenteric neurons of the colon are being generated from resident Nestin^+^ precursor cells. Next, we examined the expression of the toll-like receptors, TLR2 and TLR4. Although these have been reported to be expressed in adult enteric neurons^20, 33^, their function at this stage in life is unknown but may indicate neuronal participation in local inflammatory/immune responses^34, 35^. We found that both TLR2 and TLR4 are highly expressed by colonic ENPCs and that *in vitro* activation of TLR2 but not TLR4, induces the formation of new neurons enriched in nNOS. Prior studies have shown that TLR4 signaling may possibly regulate neuronal survival through inhibition of apoptosis, explaining the decrease in neuronal number in response to the TLR4 antagonist despite an effect on neurogenesis^15^. By contrast, inhibition of endogenous TLR2 signaling *in vivo* resulted in inhibition of nestin-derived neurogenesis in healthy mice leading to significant dysmotility and loss of colonic myenteric neurons. Participation of TLR2 in neurogenesis has also been shown in the hippocampus, consistent with our results^36^.

Because TLR2 plays a major role in gram-positive bacterial recognition via their lipoproteins^37, 38^, we examined the effects microbiota disruption by the administration of ampicillin. Profiling of the gut microbiota confirmed the disruption of natural balance of gut microbes with treatment of ampicillin, with an expected reduction of gram-positive phyla including *Firmicutes* and *Actinobacteria* and increased population of gram-negative phyla such as *Bacteroidetes*, *Proteobacteria,* and *Verrucomicrobia*. Treatment with ampicillin also impaired neurogenesis with loss of colonic neuronal number that predominantly involved nitrergic neurons, without affecting the proportion of cholinergic neurons (the remaining fraction probably represents undifferentiated neurons). Neuronal nitric oxide is a critical neurotransmitter for orderly peristalsis and in its absence, distal relaxation and bolus transit is impaired^39^. This can lead to failure of effective gastrointestinal propulsion ^6, 15^. Consistent with these mechanisms, ampicillin significantly altered the motility of the proximal colon, confirming previous reports using this drug in a cocktail of antibiotics^15, 18, 20^. The decrease in neuronal nitric oxide validates the functional importance of the change in nitrergic neuronal numbers and explains the prolonged colonic transit time. Further, discontinuation of ampicillin treatment resulted in both functional and neuronal recovery, indicating the dynamic nature of the ENS. Importantly, experiments *in vitro* and in GF mice showed no pharmacological effects of ampicillin independent of its microbial action. Finally, LTA, a cell wall constituent characteristic of gram-positive bacteria and TLR2 agonist^40^, prevented the dysbiosis-associated changes in neurogenesis and loss of global and nitrergic neurons and dysmotility, indicating the central role of gram-positive signaling in maintaining the colonic ENS. Our findings are in line with recent evidence showing that disturbing gram-positive gut microbes in neonatal period in mice using vancomycin can change the population of enteric neurons and alter gut motility, but it extends the role of gram-positive microbes into maintaining the fully developed and adult ENS^14^.

We then used germ-free mice model to further study the role of TLR2 in neurogenesis. GF mice are known to have an abnormal ENS, although the degree of abnormalities reported varies in different studies^16–18, 30^. In our study GF mice had fewer colonic neurons and a reduced proportion of nitrergic neurons, consistent with the phenotype seen with TLR2 antagonism and ampicillin-induced dysbiosis. Treatment with LTA was sufficient to increase the number of global as well as nitrergic neurons. These results are consistent with, and add to previous reports of correction of ENS abnormalities in GF mice by reconstitution of gut microbiota in the early neonatal period^18, 30^. A recent report shows an increase in the proliferation of Nestin^+^ ENPCs in the colon of GF mice as compared with conventionalized mice but speculated that the expected increase in neuronal number was not seen because of a possible compensation by increased cell loss^18^. Our results provide an alternative and more satisfying explanation. With ampicillin-induced dysbiosis, we actually found an increase in the number of Nestin^+^ cells associated with a *decrease* in neurogenesis and colonic neurons, indicating a “brake” on the maturation of neurons rather than on the proliferation of their precursors. Increased numbers of newborn neurons were found 10 days after discontinuation of ampicillin in Nestin-creER^T2^:tdTomato mice, suggesting that the putative dysbiosis-induced brake on Nestin^+^ ENPC derived neurogenesis is released, leading to restoration of neuronal numbers and colonic motility. The dysbiosis-induced brake is also released by TLR2 activation by LTA. When taken together with the results of TLR2 inactivation using a specific blocking antibody, these data indicate that TLR2 is necessary for normal neurogenesis to occur in the colon and support the therapeutic potential of TLR2 activation for selected disorders.

In conclusion, our data demonstrate for the first time the existence of neurogenesis from defined precursors in the adult colon, a process that requires gram-positive bacteria acting via a specific signaling pathway, TLR2. In its absence, ENPC continue to proliferate but cannot differentiate adequately, resulting in a loss of nitrergic colonic neurons with accompanying dysmotility. TLR2 signaling therefore appears to act at a critical check point in the differentiation of neuronal precursors. These results have important biological and clinical implications. First, further understanding of this check point in health and disease may provide important insight into the pathogenesis of acquired motility disorders. Secondly, given that antibiotic use can have long-term impact on the composition of gut microbiota even after a short exposure^13, 41, 42^, these data add ENS dysfunction to the list of important, but unintended, pathophysiological consequences of these drugs. Given the potential effects of such dysfunction on epithelial health, gut permeability, local immune responses and gut brain signaling, this paper lends more weight to the argument that such antibiotics should be used judiciously. Finally, our results suggest opportunities for the development of novel pharmacological as well as probiotic approaches directed towards specific signaling processes such as TLR2 for the treatment and/or prevention of dysbiosis-induced or other forms of colonic dysmotility such as Hirschsprung’s or other disorders.

## Abbreviations

TLR2: Toll-like Receptor 2
TLR4: Toll-like Receptor 4
ENPC: Enteric Neural Precursor Cell
ENS: Enteric Nervous System
GF: Germ-Free
LTA: Lipoteichoic Acid
LPS: Lipopolysaccharide
nNOS: neuronal Nitric Oxide Synthase
PBS: phosphate-buffered saline
LM-MP: Longitudinal muscle myenteric plexus

## Notes

**Grant support** This work was supported by National Institute of Diabetes and Digestive and Kidney Diseases Grant R01DK080920 (to P.J.P.); Bloomberg Philanthropies (to C.L.S); Grant P30 DK089502 (to S.S.Y); and a grant from the Amos family (to P.J.P).

**Disclosures:** The authors have declared that no conflict of interest exists.

## References

1. Furness JB. The enteric nervous system and neurogastroenterology. Nat Rev Gastroenterol Hepatol 2012;9:286–94.

2. Gariepy CE. Intestinal motility disorders and development of the enteric nervous system. Pediatr Res 2001;49:605–13.

3. De Giorgio R, Camilleri M. Human enteric neuropathies: morphology and molecular pathology. Neurogastroenterol Motil 2004;16:515–31.

4. Joseph NM, He S, Quintana E, et al. Enteric glia are multipotent in culture but primarily form glia in the adult rodent gut. J Clin Invest 2011;121:3398–411.

5. Kugler EM, Michel K, Zeller F, et al. Mechanical stress activates neurites and somata of myenteric neurons. Front Cell Neurosci 2015;9:342.

6. Anitha M, Reichardt F, Tabatabavakili S, et al. Intestinal dysbiosis contributes to the delayed gastrointestinal transit in high-fat diet fed mice. Cell Mol Gastroenterol Hepatol 2016;2:328–339.

7. Neunlist M, Schemann M. Nutrient-induced changes in the phenotype and function of the enteric nervous system. J Physiol 2014;592:2959–65.

8. Wafai L, Taher M, Jovanovska V, et al. Effects of oxaliplatin on mouse myenteric neurons and colonic motility. Front Neurosci 2013;7:30.

9. Bagyanszki M, Bodi N. Gut region-dependent alterations of nitrergic myenteric neurons after chronic alcohol consumption. World J Gastrointest Pathophysiol 2015;6:51–7.

10. Cotillard A, Kennedy SP, Kong LC, et al. Dietary intervention impact on gut microbial gene richness. Nature 2013;500:585–8.

11. Claesson MJ, Jeffery IB, Conde S, et al. Gut microbiota composition correlates with diet and health in the elderly. Nature 2012;488:178–84.

12. Modi SR, Collins JJ, Relman DA. Antibiotics and the gut microbiota. J Clin Invest 2014;124:4212–8.

13. Dethlefsen L, Huse S, Sogin ML, et al. The pervasive effects of an antibiotic on the human gut microbiota, as revealed by deep 16S rRNA sequencing. PLoS Biol 2008;6:e280.

14. Hung LY, Boonma P, Unterweger P, et al. Neonatal Antibiotics Disrupt Motility and Enteric Neural Circuits in Mouse Colon. Cell Mol Gastroenterol Hepatol 2019.

15. Anitha M, Vijay-Kumar M, Sitaraman SV, et al. Gut microbial products regulate murine gastrointestinal motility via Toll-like receptor 4 signaling. Gastroenterology 2012;143:1006–16 e4.

16. McVey Neufeld KA, Mao YK, Bienenstock J, et al. The microbiome is essential for normal gut intrinsic primary afferent neuron excitability in the mouse. Neurogastroenterol Motil 2013;25:183–e88.

17. McVey Neufeld KA, Perez-Burgos A, Mao YK, et al. The gut microbiome restores intrinsic and extrinsic nerve function in germ-free mice accompanied by changes in calbindin. Neurogastroenterol Motil 2015;27:627–36.

18. De Vadder F, Grasset E, Manneras Holm L, et al. Gut microbiota regulates maturation of the adult enteric nervous system via enteric serotonin networks. Proc Natl Acad Sci U S A 2018.

19. Della Mina E, Borghesi A, Zhou H, et al. Inherited human IRAK-1 deficiency selectively impairs TLR signaling in fibroblasts. Proc Natl Acad Sci U S A 2017;114:E514–E523.

20. Brun P, Giron MC, Qesari M, et al. Toll-Like Receptor 2 Regulates Intestinal Inflammation by Controlling Integrity of the Enteric Nervous System. Gastroenterology 2013;145:1323–1333.

21. Kulkarni S, Micci MA, Leser J, et al. Adult enteric nervous system in health is maintained by a dynamic balance between neuronal apoptosis and neurogenesis. Proc Natl Acad Sci U S A 2017.

22. Chandrasekharan BP, Kolachala VL, Dalmasso G, et al. Adenosine 2B receptors (A(2B)AR) on enteric neurons regulate murine distal colonic motility. FASEB journal: official publication of the Federation of American Societies for Experimental Biology 2009;23:2727–34.

23. Becker L, Nguyen L, Gill J, et al. Age-dependent shift in macrophage polarisation causes inflammation-mediated degeneration of enteric nervous system. Gut 2018;67:827–836.

24. Shirasawa S, Yunker AM, Roth KA, et al. Enx (Hox11L1)-deficient mice develop myenteric neuronal hyperplasia and megacolon. Nat Med 1997;3:646–50.

25. Pigott B, Bartus K, Garthwaite J. On the selectivity of neuronal NOS inhibitors. Br J Pharmacol 2013;168:1255–65.

26. Korenaga K, Micci MA, Taglialatela G, et al. Suppression of nNOS expression in rat enteric neurones by the receptor for advanced glycation end-products. Neurogastroenterol Motil 2006;18:392–400.

27. Dwyer MA, Bredt DS, Snyder SH. Nitric oxide synthase: irreversible inhibition by L-NG-nitroarginine in brain in vitro and in vivo. Biochem Biophys Res Commun 1991;176:1136–41.

28. Hunt D, Drake LA, Drake JR. Murine macrophage TLR2-FcgammaR synergy via FcgammaR licensing of IL-6 cytokine mRNA ribosome binding and translation. PLoS One 2018;13:e0200764.

29. Neal MD, Jia H, Eyer B, et al. Discovery and validation of a new class of small molecule Toll-like receptor 4 (TLR4) inhibitors. PLoS One 2013;8:e65779.

30. Collins J, Borojevic R, Verdu EF, et al. Intestinal microbiota influence the early postnatal development of the enteric nervous system. Neurogastroenterol Motil 2014;26:98–107.

31. Muller PA, Koscso B, Rajani GM, et al. Crosstalk between Muscularis Macrophages and Enteric Neurons Regulates Gastrointestinal Motility. Cell 2014;158:1210.

32. Shen L, Weber CR, Raleigh DR, et al. Tight junction pore and leak pathways: a dynamic duo. Annu Rev Physiol 2011;73:283–309.

33. Rumio C, Besusso D, Arnaboldi F, et al. Activation of smooth muscle and myenteric plexus cells of jejunum via Toll-like receptor 4. J Cell Physiol 2006;208:47–54.

34. Coquenlorge S, Duchalais E, Chevalier J, et al. Modulation of lipopolysaccharide-induced neuronal response by activation of the enteric nervous system. J Neuroinflammation 2014;11:202.

35. Burgueno JF, Barba A, Eyre E, et al. TLR2 and TLR9 modulate enteric nervous system inflammatory responses to lipopolysaccharide. J Neuroinflammation 2016;13:187.

36. Rolls A, Shechter R, London A, et al. Toll-like receptors modulate adult hippocampal neurogenesis. Nat Cell Biol 2007;9:1081–8.

37. Takeuchi O, Hoshino K, Kawai T, et al. Differential roles of TLR2 and TLR4 in recognition of gram-negative and gram-positive bacterial cell wall components. Immunity 1999;11:443–51.

38. Chandra R, Mellis B, Garza K, et al. Remnant lipoprotein size distribution profiling via dynamic light scattering analysis. Clin Chim Acta 2016;462:6–14.

39. Mizuta Y, Takahashi T, Owyang C. Nitrergic regulation of colonic transit in rats. Am J Physiol 1999;277:G275–9.

40. Triantafilou M, Manukyan M, Mackie A, et al. Lipoteichoic acid and toll-like receptor 2 internalization and targeting to the Golgi are lipid raft-dependent. J Biol Chem 2004;279:40882–9.

41. Sjolund M, Wreiber K, Andersson DI, et al. Long-term persistence of resistant Enterococcus species after antibiotics to eradicate Helicobacter pylori. Ann Intern Med 2003;139:483–7.

42. Yassour M, Vatanen T, Siljander H, et al. Natural history of the infant gut microbiome and impact of antibiotic treatment on bacterial strain diversity and stability. Sci Transl Med 2016;8:343ra81.

