## Supplementary material for "Intestinal Bacteria Maintain Adult Enteric Nervous System and Nitrergic Neurons via Toll-like Receptor 2-induced Neurogenesis in Mice": SI material

**Supplementary Data**

**Animals.**

Nestin-GFP mice. Adult male Nestin-GFP mice in a C57/BL background^1^ were used for imaging of the Nestin-GFP network in different cohorts of antibiotic treated and vehicle groups and FACs analysis for expression of TLR2 by Nestin ENPCs.

Nestin-creER^T2^:tdTomato mice. Nestin-creER^T2^ mice (Jaxmice #016261) were bred with floxed tdTomato (Ai14 mouse; Jaxmice #007908) mice to create Nestin-creER^T2^:tdTomato mice. For the *in vivo* experiments with Nestin-creER^T2^, only mice that were homozygous for tdTomato were used.

ChAT-cre:tdTomato mice. ChAT-cre mice (Jaxmice #018957) were bred with floxed tdTomato (Ai14) mice to generate ChAT-cre:tdTomato mice.

Germ-free (GF) mice. Germfree mice were maintained in the Gnotobiotic Core of the Center for Gastrointestinal Biology and Disease at Johns Hopkins. Gnotobiotic animal use protocols were approved by the Institutional Animal Care and Use Committee, Johns Hopkins University.

**Bead latency test.**

Distal colonic transit time was measured at 1, 2, 3, and 4 weeks after treatment. A 3-mm glass bead was placed 2 cm proximal to the anal opening using a plastic Pasteur pipette lightly lubricated with lubricating jelly. Distal colonic transit time was assessed by measuring the amount of time between bead placement and expulsion of the bead ^2^.

**16S rRNA gene PCR amplicon sequencing and library preparation.** Extracted DNA was normalized so that 3 μL of 3 ng/μL was input into each 25 μL PCR reaction. Primers and reaction conditions used in the study were described previously ^3^. Briefly, 3 μL of 5 μM primers were added to 19 μL Accuprime Pfx Supermix (ThermoFisher, Waltham, MA). Samples were denatured for 2 min at 95 °C, and then 25 cycles of 95 °C for 20 seconds, 55 °C for 15 seconds, and 72 °C for 2 min. Final extension was at 72 °C for 10 minutes. Following completion of PCR reaction, all samples were pooled in equal volumes and purified with Ampure XP beads (Beckman Coulter, Indianapolis, IN) according to manufacturer’s specifications. The purified library was quantified, loaded, and sequenced in a 600 cycle run on an Illumina MiSeq using V3 chemistry according to manufacturer's specifications (Illumina, San Diego, CA) and protocols described previously ^3^. Details of 16S rRNA gene amplicon sequence preprocessing and denoising are in supplementary data.

**Tissue preparation and cell isolation.** Longitudinal muscle myenteric plexus (LM-MP) tissue was isolated as described before ^4^. Briefly, mice were anesthetized with isoflurane and sacrificed by cervical dislocation. A laparotomy was performed, the colon was removed, and the luminal contents were flushed with PBS containing penicillin-streptomycin (PS; Invitrogen), then cut into 2-cm-long segments and further lavaged in PBS containing PS. A superficial longitudinal incision was made along the serosal surface and the myenteric plexus preparations was peeled off from the underlying tissue using a wet sterile cotton swab^4, 5^ and placed in Opti-MEM medium (Invitrogen) containing PenStrep (Invitrogen). Myenteric plexus was dissociated in a digestion buffer consisting of M199 media (Invitrogen) containing 0.1% BSA, 1 mM CaCl2, 20 mM Hepes, 150 μM P188, 50 U/mL DNase I (Worthington), and 1.1 mg/mL collagenase (Sigma) for 40 min at 37 °C and 5% CO2. The tissue was washed in PBS with 1% BSA then first passed through a 70-μm and subsequently through a 40-μm nylon mesh cell strainer to yield single cells.

**Microbiome statistical analyses.** All microbiome statistical analyses were performed in Rv. 3.3.3 (R Core Team, 2017), and we attributed statistical significance at p < 0.05. Data were checked for normality and homogeneity of variance and data that could not be transformed to attain normality were analyzed using non-parametric tests. To determine if overall bacterial community composition was affected by ampicillin treatment, we used multivariate nonparametric ANOVA of dissimilarities (PERMANOVA) with 999 permutations, using the Adonis function in the Vegan package in R based on Bray-Curtis distance ^6^. Principle coordinate analysis (PCoA) was performed on the microbial communities to visualize differences between groups ^7^. To determine alpha diversity, we calculated the Shannon-Wiener index and used a linear model to determine statistically significant changes in the alpha diversity ^8, 9^. Finally, linear models were performed to compare the effects of ampicillin on bacterial phyla and families present in the microbiome immediately following 2 weeks of ampicillin and 10 days after discontinuation of ampicillin.

**Examination of Nestin derived neurogenesis in the colon.**

*Anatomical Location of Nestin^+^ Cells in the Adult Colon.* To examine if Nestin^+^ cells are present in the colonic myenteric plexus, we used a Nestin-GFP reporter mice, as previously reported in other works from our laboratory. The colonic segments were harvested and flushed by PBS to remove contents. After, the tissue was fixed in 4% PFA overnight, myenteric plexus was isolated and immunostained for HuC/D and counterstained with DAPI as explained above. Slides were imaged by laser confocal microscopy using a Zeiss LSM 510 META.

*Examination of neurogenesis from Nestin^+^ cells.* To study whether new colonic myenteric neurons are formed from Nestin^+^ cells, we used Nestin-creER^T2^:tdTomato mice as a model. In these mice, Nestin^+^ cells and their derivative cells express tdTomato after induction with tamoxifen. Mice were sacrificed 6 days after tamoxifen induction and their colonic myenteric plexus were isolated, fixed and immune-stained with anti-sera against HuC/D as described above. Number of neurons that express tdTomato was enumerated.

**Role of TLR2 in neurogenesis from Nestin^+^ cells.**

*Examining the expression of TLR2 and TLR4 by Nestin^+^ cells and their derivative neurons.* We first examined if Nestin-GFP^+^ cells express these receptors by using Nestin-GFP reporter mice. Mice were sacrificed, and their colonic myenteric plexus were isolated as described before, myenteric plexus was then dissociated in a digestion buffer consisting of M199 media (Invitrogen) containing 0.1% BSA, 1 mM CaCl2, 20 mM Hepes, 150 μM P188, 50 U/mL DNase I (Worthington), and 1.1 mg/mL collagenase (Sigma) for 40 min at 37 °C and 5% CO2. The tissue was washed in PBS with 1% BSA then first passed through a 70-μm and subsequently through a 40-μm nylon mesh cell strainer to yield single cells. Flow analyses to study the expression of TLR2 and TLR4 in Nestin-GFP^+^ cells isolated from myenteric plexus of Nestin-GFP mouse, was performed on BD LSR II.UV after staining with respective antibodies (Table S1).

To study the expression of TLR2 and TLR4 by enteric neurons, we used Nestin-creER^T2^:tdTomato mice. After induction with tamoxifen as described above, mice were sacrificed, and myenteric plexus were isolated and dissociated in a digestion buffer as described above. Cells obtained from myenteric plexus were then plated and cultured for 14 days. After 14 days, cells were fixed in 2% PFA for 15 minutes at 22°C, washed, treated with permeabilization buffer (5% NGS and 0.3% Triton X) then stained with antibody against TLR2 and TLRR2 separately (Table S1).

**Examination of TLR2 and TLR4 inhibition effects on ENS.**

Nestin-creER^T2^:tdTomato animals received an IP dose of T2.5 (anti-TLR2 monoclonal antibody; 1mg/mouse; Abeomics, San Diego, CA) immediately after they were induced with tamoxifen. The myenteric plexus from cecum of these mice were isolated 14 days after tamoxifen induction, fixed and fixed and immunostained for HuC/D and nNOS and counterstained with DAPI. Slides were image using a Leica 710 confocal microscope. The images then were analyzed using Fiji software (fiji.sc).

To study the role of TLR4 in modulating neurogenesis, TLR4 blocker C34 was administered to the Nestin-creER^T2^:tdTomato mice via daily oral gavage at a concentration of 1 mg/kg at the same time as tamoxifen induction and continued for 14 days. After 14 days, myenteric plexus from cecum was isolated and immunostained for HuC/D. Slides were image using a Leica 710 confocal microscope. The images then were analyzed using Fiji software (fiji.sc).

**Examination of reversibility of ampicillin effect.**

To study the reversibility of the effects of ampicillin on gut motility and enteric nervous system, 8-12-week old adult C57/BL male mice (n=5/group) were treated with ampicillin (0.5 gram/L) through drinking water containing 4 g/L Splenda in addition to the control group for 4 weeks. Ampicillin was then discontinued and WGTT was measured at day 0, 3, 5 and 10 after discontinuation of ampicillin, using carmine dye protocol as described above. Mice were sacrificed after 10 days and their colonic myenteric plexus were isolated and fixed and immunostained for HuC/D and counterstained with DAPI as described above. Slides were imaged using a Leica 510 confocal microscope. The images then were analyzed using Fiji software (fiji.sc). ANOVA was used to compare the WGTT and neuronal counts and ganglion density per 10X high power field between different groups.

**Examining the effects of ampicillin treatment on Nestin^+^ enteric neural precursor cells and neurogenesis.**

*Examination of the effect of Ampicillin on Nestin+ cells.* To study the impact of Ampicillin treatment on Nestin-GFP^+^ cells, we used Nestin-GFP reporter mice and treated them with Ampicillin 0.5 gram/l for 2 weeks. Mice were then sacrificed, and the colonic myenteric plexus were isolated, fixed and stained with HuC/D and counterstained with DAPI as described above.

*Examination of the effect of Ampicillin on neurogenesis.* Nestin-creER^T2^:tdTomato animals were induced with tamoxifen. Mice were then started on ampicillin as explained above. Cohorts of animals were then sacrificed after 2 weeks of ampicillin, 10 days after ampicillin discontinuation, or 2 weeks of placebo. The myenteric plexus from cecum of these mice were isolated 14 days after tamoxifen induction, fixed and immunostained for HuC/D and nNOS and counterstained with DAPI. Slides were imaged using a Leica 710 confocal microscope. The images then were analyzed using Fiji software (fiji.sc).

**Examining the effects of lipoteichoic acid (LTA) on ampicillin-induced changes in gut motility and enteric nervous system (ENS).**

To further study the role of gut microbiota in modulating the effects of ampicillin on gut motility and ENS, we used lipoteichoic acid (LTA), which is the bacterial wall product of gram-positive bacteria. Adult C57BL/6 mice were treated with Ampicillin+Saline (0.5 gram/L), Ampicillin+LTA (Lipoteichoic Acid, 2.5 mg/Kg through oral gavage) or control for 2 weeks. WGTT was measured using the carmine dye protocol as described above. Mice were then sacrificed and their colonic myenteric plexus were isolated, fixed and stained with antisera against HuC/D,and nNOS with the same method described earlier.

**Experiments in Germ-Free Mice.**

*To examine the effects of LTA on enteric nervous system.* C57BL/6 GF mice were given LTA (5 mg/liter) through drinking water for different durations of 2 and 3 weeks. The myenteric plexus from cecum of these mice were isolated after 14 days, fixed and immunostained for HuC/D and nNOS and counterstained with DAPI. Slides were imaged using a Leica 710 confocal microscope. The images then were analyzed using Fiji software (fiji.sc).

*To examine whether ampicillin has direct pharmacologic effect on GI motility and colonic enteric neurons*. We studied the effects of Ampicillin in germ-free mice. GF mice were treated with ampicillin or Splenda through drinking water for 2 weeks. WGTT was measured at 2 weeks using the carmine dye protocol described earlier. Mice were then sacrificed, and their colonic myenteric plexus were isolated, fixed and immune-stained with antisera against HuC/D and nNOS.

**16S rRNA gene amplicon sequence preprocessing.**

All samples were sequenced together on the same run of the sequencing instrument, and demultiplexed according to sample-specific barcodes as described in Kozich et al. Read quality control and the resolution of amplicon sequence variants (ASV) were completed using the DADA2 R package (v1.7.5) ^10^. Per dada2 authors’ recommendations, all sequences were trimmed at fixed positions following inspection of aggregate positional sequence quality in order to omit severe drops in quality due to end effects (Figure S7). Forward reads were trimmed before position 10 and after position 230 (5’ and 3’ ends, respectively). The reverse reads were not used due to low quality. Filters were applied to remove individual reads from the dataset if they contained any Ns, mapped to any part of the phiX genome, or were predicted based on quality values to have a maximum expected error greater than 2 (‘filterAndTrim’ function).

**16S rRNA gene amplicon sequence denoising.**

Error model estimation was calculated on a randomly-selected subset of 30 samples out of 225 total from the run using the ‘learnErrors’ function. This aggregate model was applied to every sample during final independent amplicon sequence denoising via the ‘dada’ function, with other parameters set to their default values. Chimera detection and removal was performed via the `removeBimeraDenovo` function (308 out of 2529 denoised sequences, ~12.2%). The taxonomy of each amplicon sequence was classified independently via SINTAX using the RDP v.16 of the 16S rRNA gene sequence database as formatted and distributed on the USEARCH website ^11^. ASVs were aligned using the R MSA package and a maximum likelihood phylogenetic tree was computed using the GTR model with optimization of the proportion of invariable sites and gamma rate parameter, via the phangorn package, as described in a recently published workflow ^12-14^. The resulting tree was combined with the table of ASV counts for each sample and merged with experimental design data for loading into the phyloseq package ^15^.

### Microbiome sample diversity and differential abundance comparisons. The phyloseq package was used to compute the Shannon Index of each sample as a measure of alpha diversity. The phyloseq package was also used to compute the pairwise weighted-UniFrac distance matrix (‘distance’ function), which was then decomposed via Principal Coordinate Analysis (“PCoA”, ‘ordinate’ function).

Differential abundance of ASVs between conditions was conducted via a paired Wilcoxon test on the centered log-ratio (CLR) transformed values of a set of Monte-Carlo Dirichlet resampling instances, as implemented in the ALDEx2 Bioconductor/R package ^16^. A multiple testing correction was applied to the statistical significance of each result (False Discovery Rate, FDR).

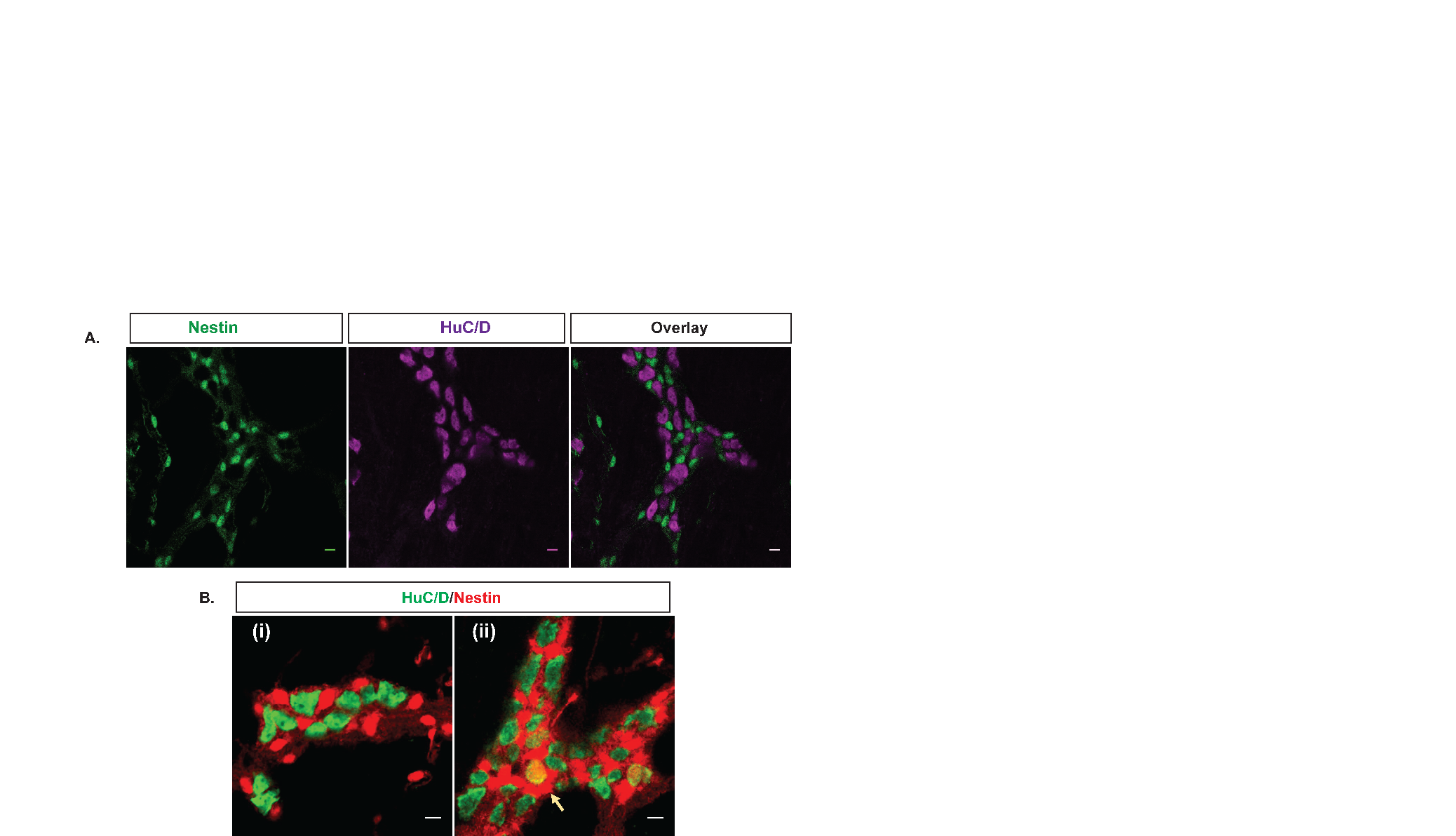

**Figure S1. Nestin^+^ cells exist in the colonic myenteric plexus and generate neurons in vivo.** A panel are representative photomicrographs of Nestin-GFP mice after immunostaining with the neuronal marker HuC/D. B (i-ii) are representative photomicrographs of colonic myenteric plexus ganglion of Nestin-creER^T2^:tdTomato mice after tamoxifen induction and immunostaining with the neuronal marker HuC/D. B (i). 12 hours after tamoxifen induction, there are no neurons that express tdTomato. B (ii). 6 days after tamoxifen induction, neurons are seen expressing tdTomato (arrow) indicating an origin from Nestin^+^ cells. Scale bar is 10 µm.

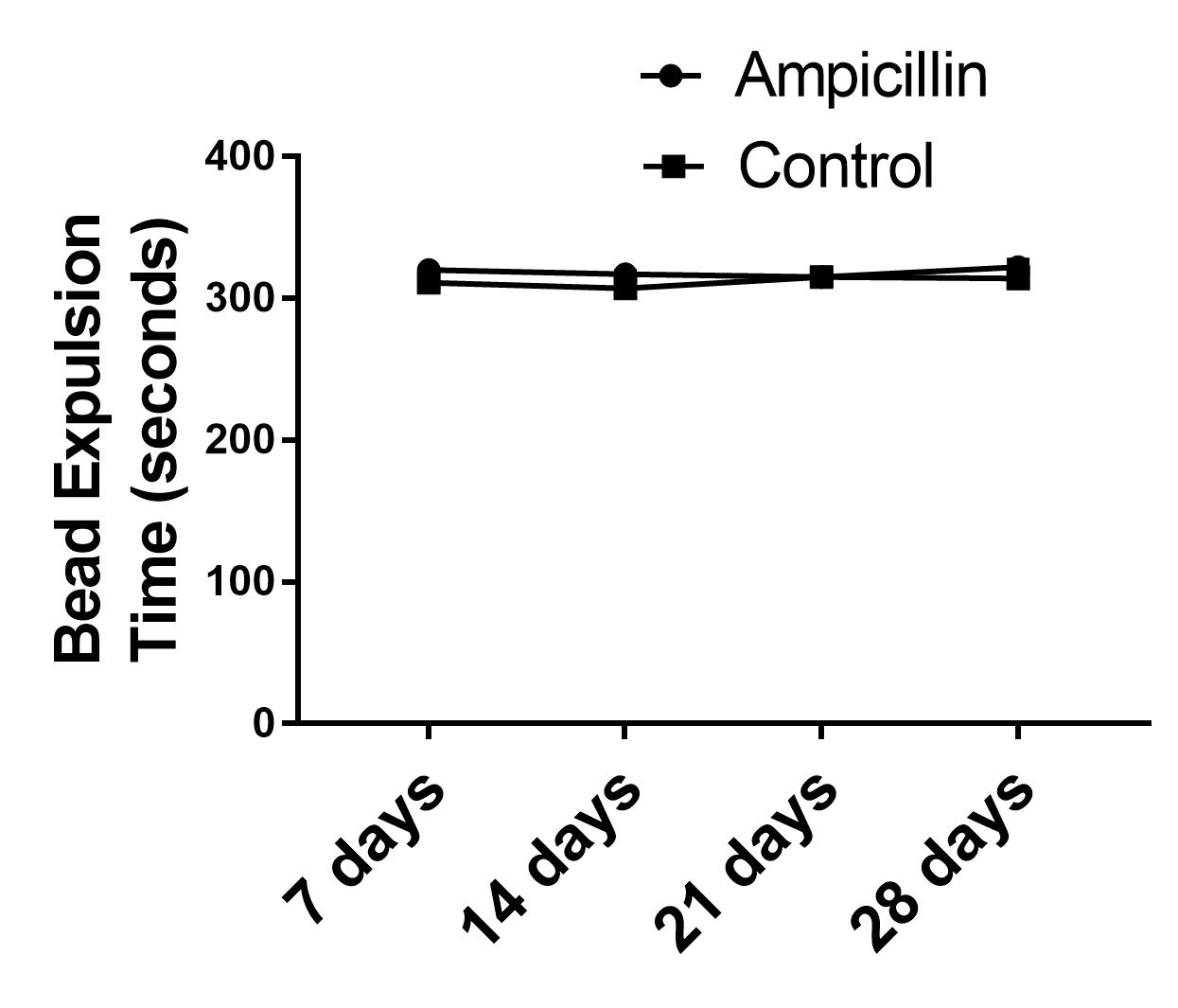

**Figure S3. Ampicillin treatment did not affect bead expulsion latency test, a measure of distal colonic transit time.** Distal colonic transit time was similar to the control group after 1, 2, 3, and 4 weeks of treatment with ampicillin.

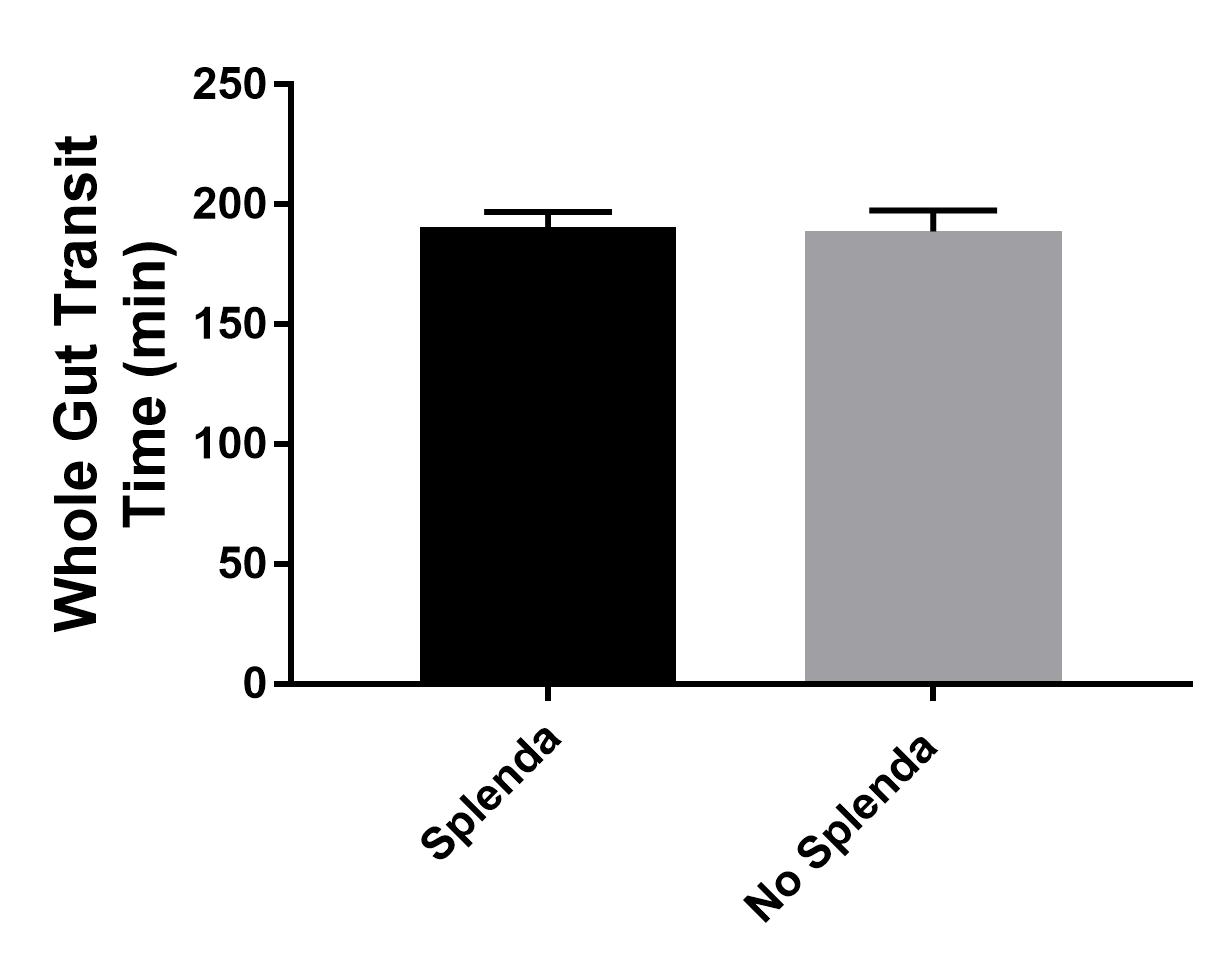

**Figure S2. Splenda through drinking water does not affect whole gut transit time.** Grouped results of mean whole gut transit time in two different groups of Splenda and no Splenda (p=0.87).

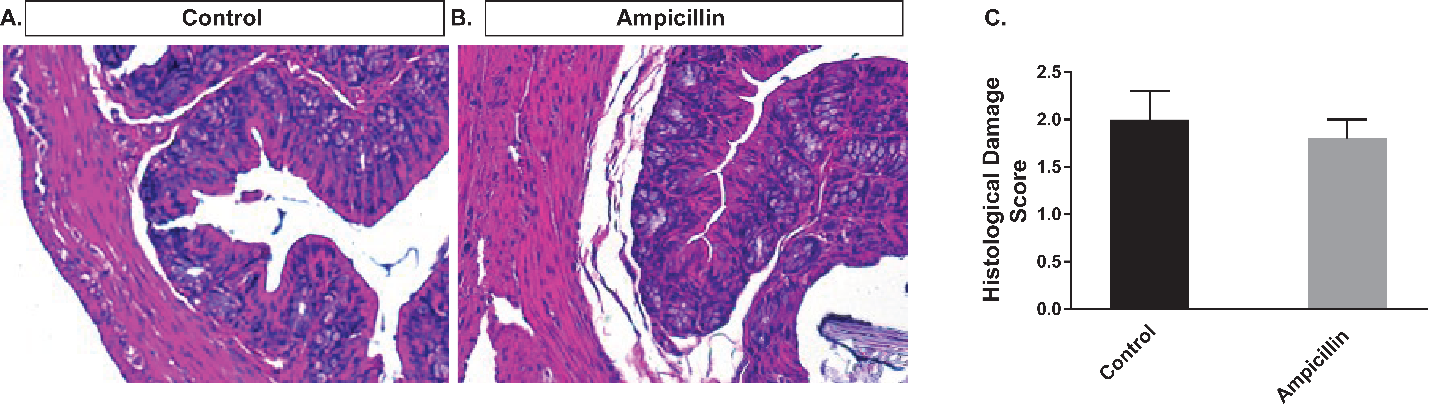

**Figure S4. Ampicillin treatment does not result in epithelial inflammation or injury, as assessed by histological damage score calculation.** (A-B) Representative hematoxylin and eosin (H&E) staining of the colon cross section in vehicle (A) and ampicillin treated (B) wild type mice (magnification 40x). C. Grouped results of histological damage score calculation (p=NS).

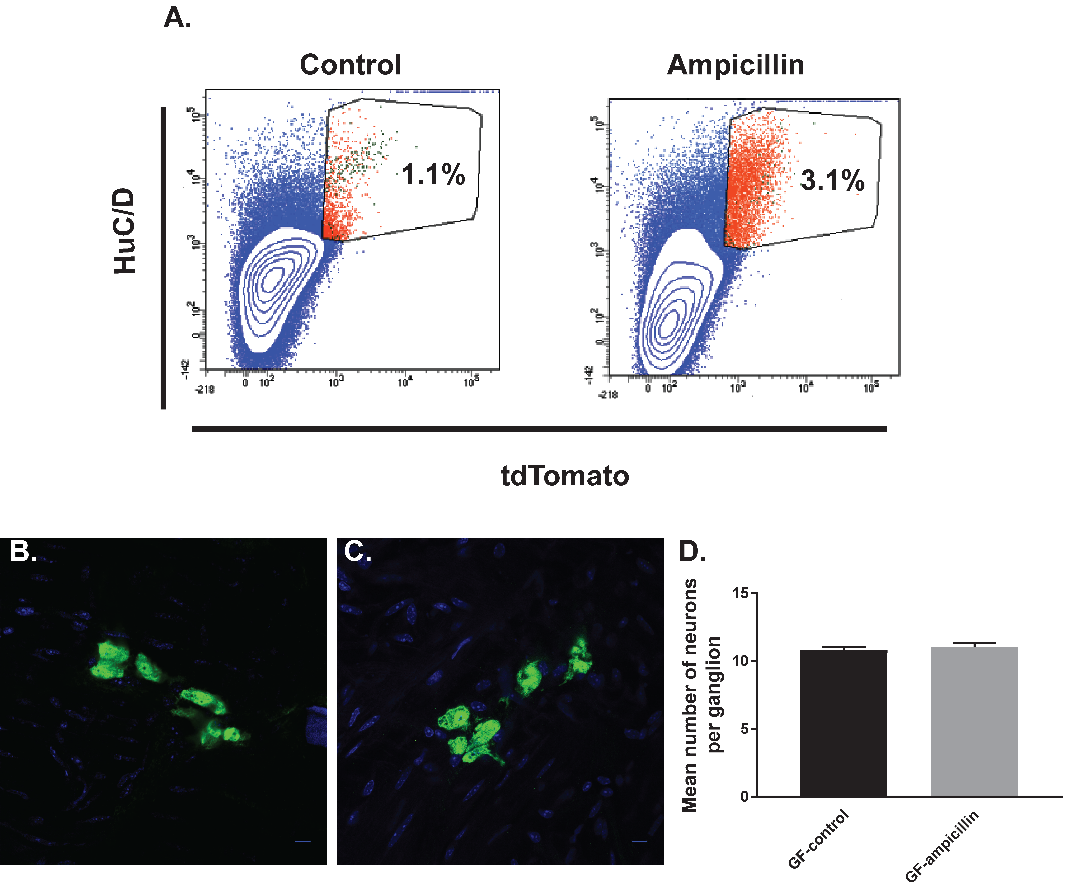

**Figure S5. Ampicillin does not have a direct toxic effect on the enteric neurons in vitro or in vivo.** (A) Pseudocolor plots of flow cytometry data of cells isolated from colonic myenteric plexus of Nestin-creERT2:tdTomato mice after tamoxifen induction and being cultured for 14 days and stained for HuC/D. (A. left panel) Flow cytometry from control culture group showing newly formed neurons from Nestin^+^ ENPCs that express tdTomato (top right quadrant). (A. right panel) Addition of ampicillin did not increase the number of newly formed neurons. B-C are representative photomicrographs of colonic myenteric plexus after immunostaining with the neuronal marker HuC/D (green); nuclei are stained with DAPI (blue). (B) Immunostaining with neuronal marker HuC/D (green) shows baseline myenteric ganglion density in GF colon, that (C) was not altered with ampicillin treatment in GF colon. (D) Grouped results of neuronal numbers per ganglia under various conditions. Scale bar is 10 µm in B-C.

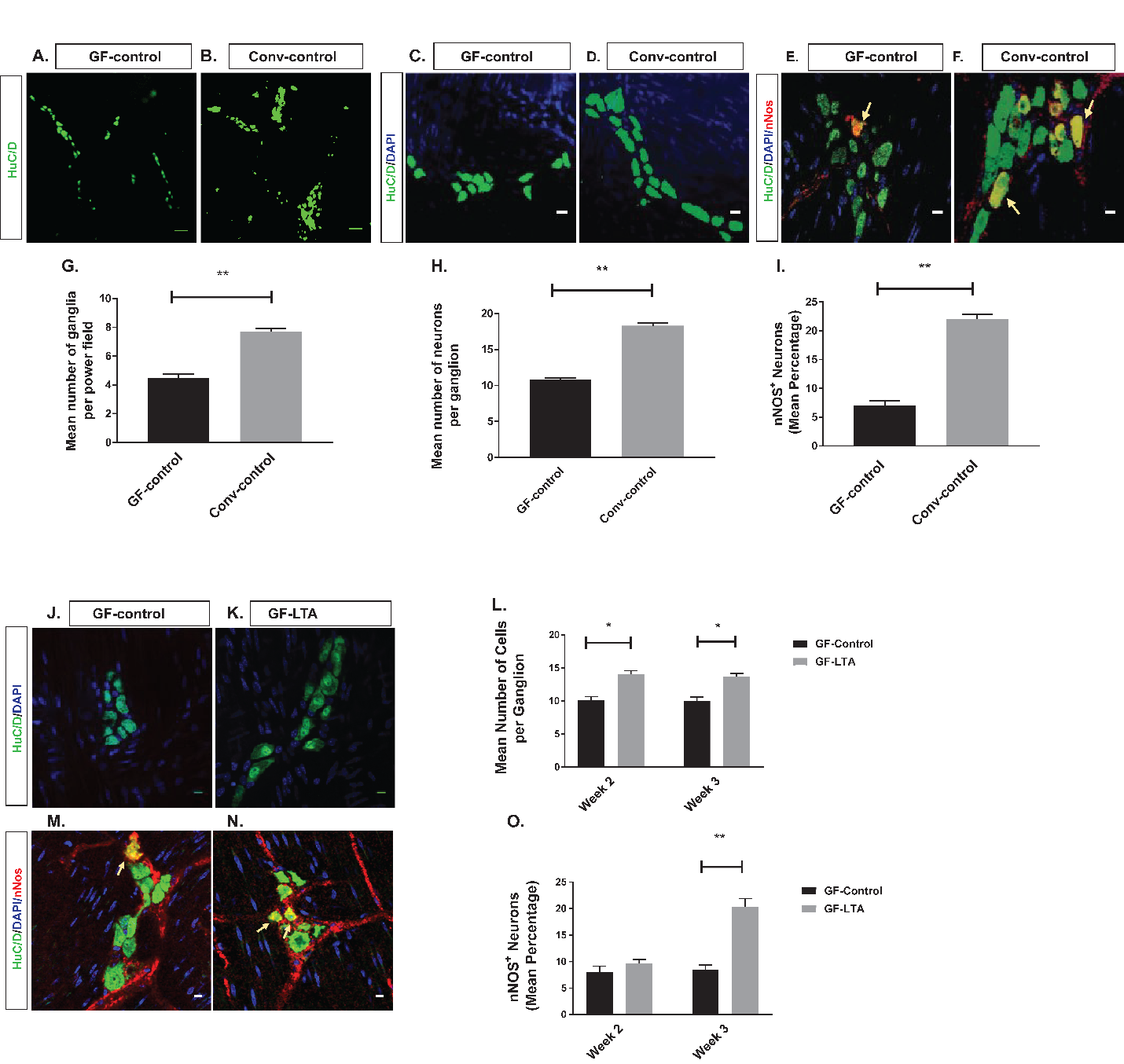

**Figure S6. TLR2 agonist, LTA increases restores neuronal number in germ free (GF) mice.** A-D and J-K are representative photomicrographs of colonic myenteric plexus after immunostaining with the neuronal marker HuC/D (green); nuclei are stained with DAPI (blue). E-F and M-N are representative photomicrographs of a colonic myenteric ganglion after immunostaining with the neuronal marker HuC/D (green) and nitrergic neuron marker nNOS (red); nuclei are stained with DAPI (blue). (A) Immunostaining with neuronal marker HuC/D (green) shows baseline myenteric ganglion density in GF colon. (B) Shows baseline myenteric ganglionic density in colons from adult conventional mice. (C) Immunostaining with neuronal marker (HuC/D) shows neurons in a representative colonic myenteric ganglion in GF mice. (D) Baseline number of neurons per ganglion in conventional mice. (E) GF mice had reduced percentage of nitrergic neurons (arrow). (F) Conventional mice had significantly higher percentage of nitrergic neurons (arrows) per ganglion. (G) Grouped results of ganglion density under various conditions (**p<0.01). (H) Grouped results of neuronal numbers per ganglia under various conditions (**p<0.01). (I) Grouped results of percentage of nitrergic neurons per ganglion under various conditions (**p<0.01). (J) Immunostaining with antisera against HuC/D (green) of myenteric plexus in GF-control mice at 3 weeks (K) 3 weeks of LTA treatment increased the number of neurons per ganglion. (L) Grouped results of neuronal number per ganglion under various conditions (*p<0.05). (M) Representative image of a normal myenteric ganglion including nitrergic neurons (arrow). (N) 3 weeks of LTA treatment increased the number and percentage of nitrergic neurons (arrows). (O) Grouped results of percentage of nitrergic neurons per ganglion under various conditions (**p<0.01). Scale bar is 10 µm.

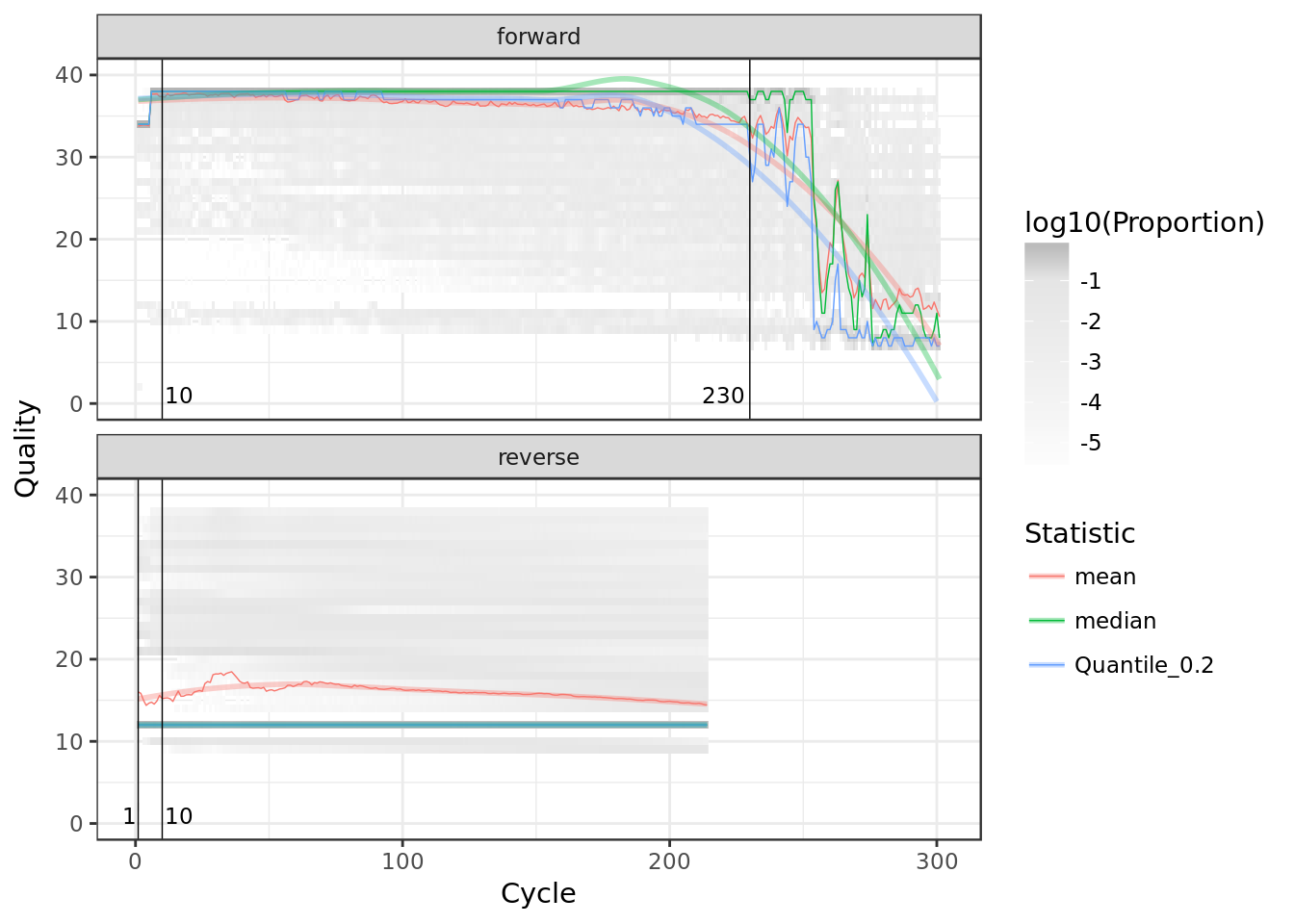

**Figure S7. Aggregate positional quality of 16S rRNA amplicon sequence reads.** Forward and reverse reads are separated into top and bottom panels, respectively. The left and right trimming positions for the forward read are indicated with a vertical black line.

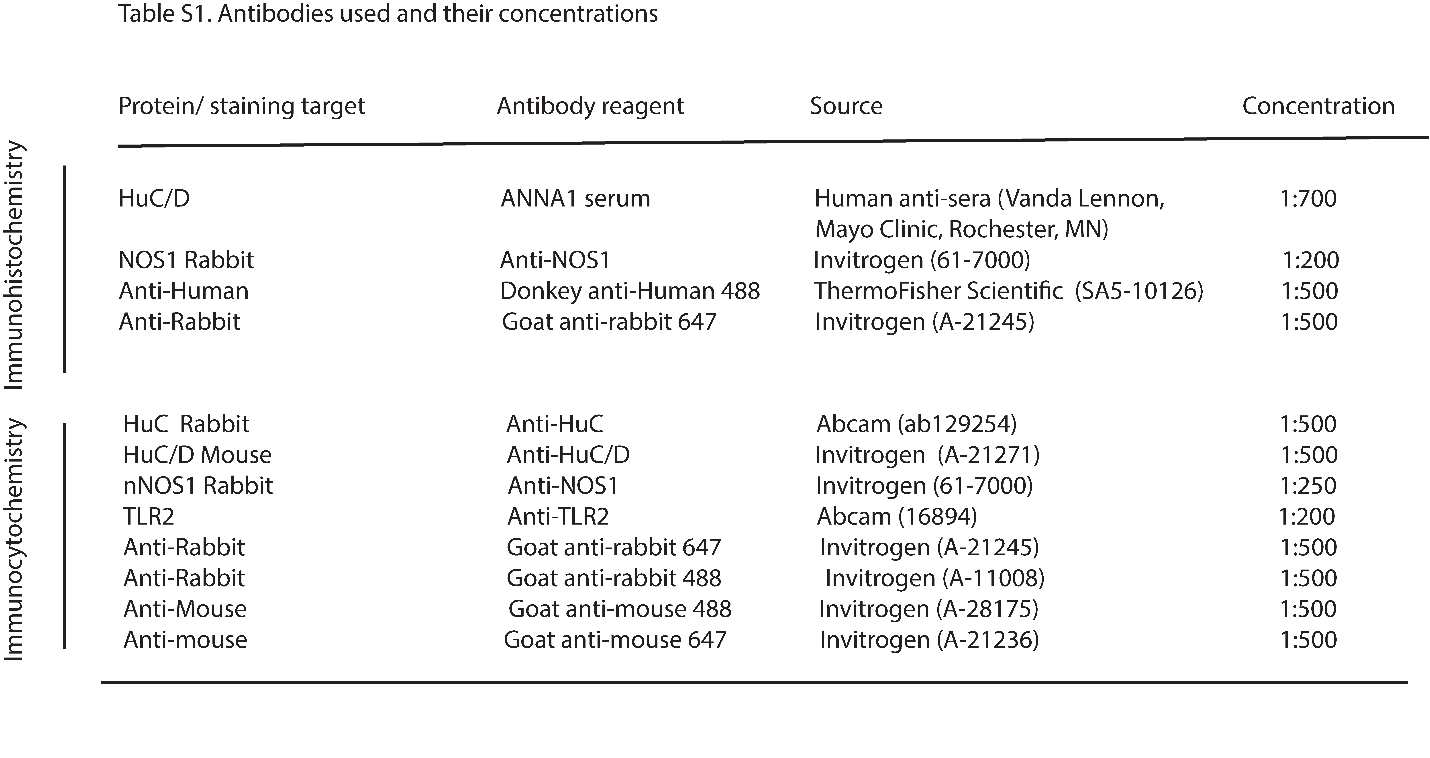

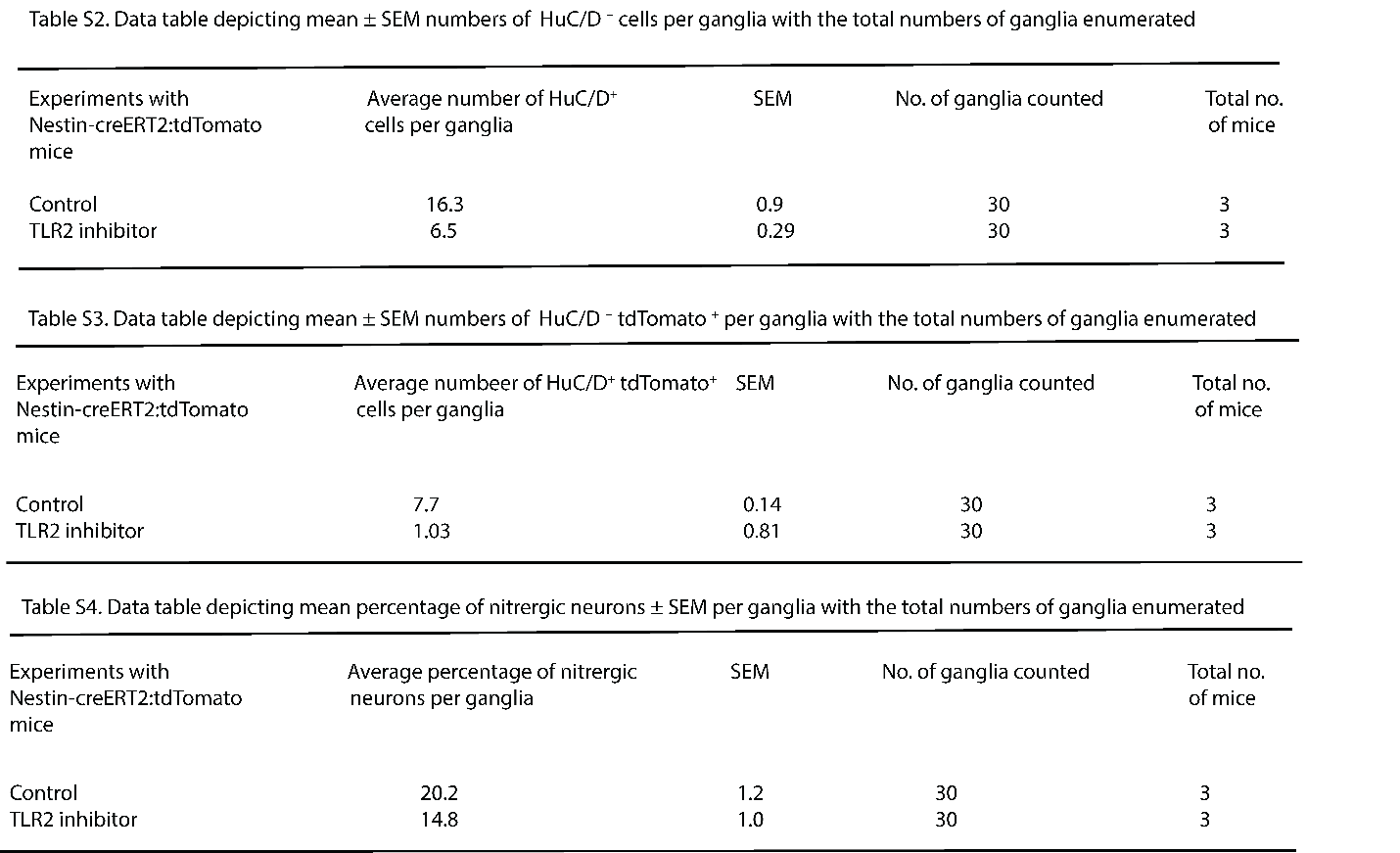

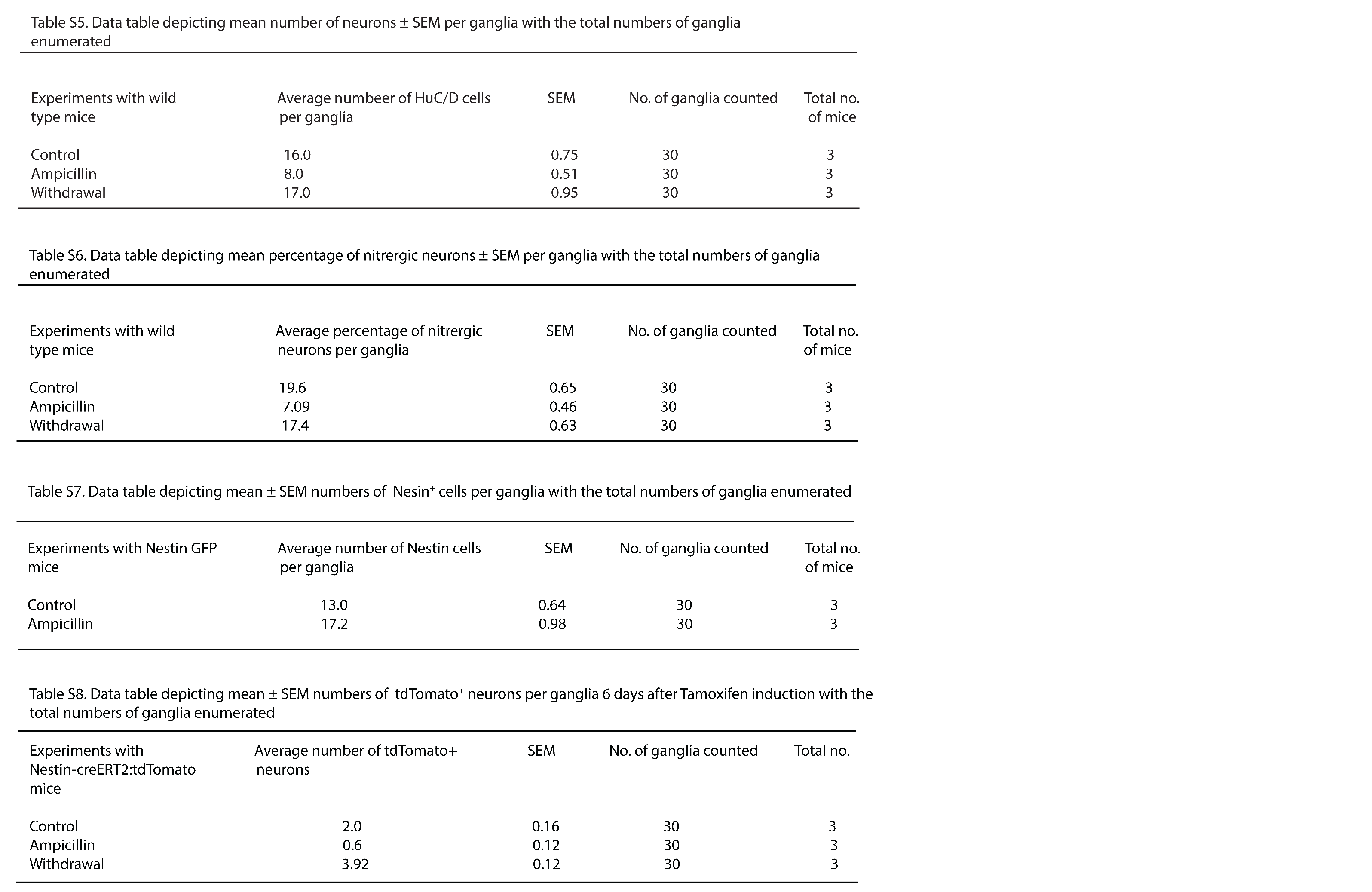

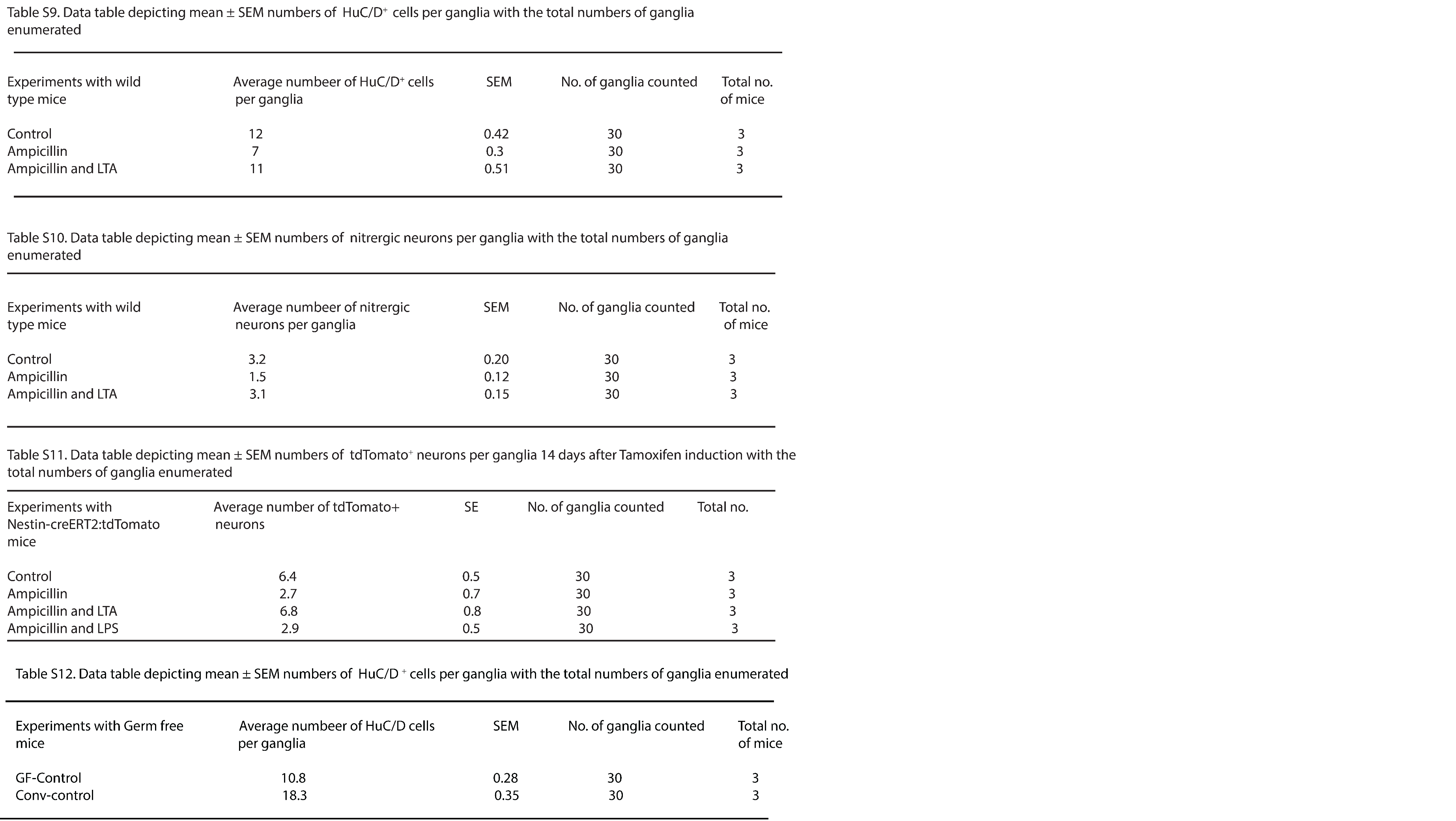

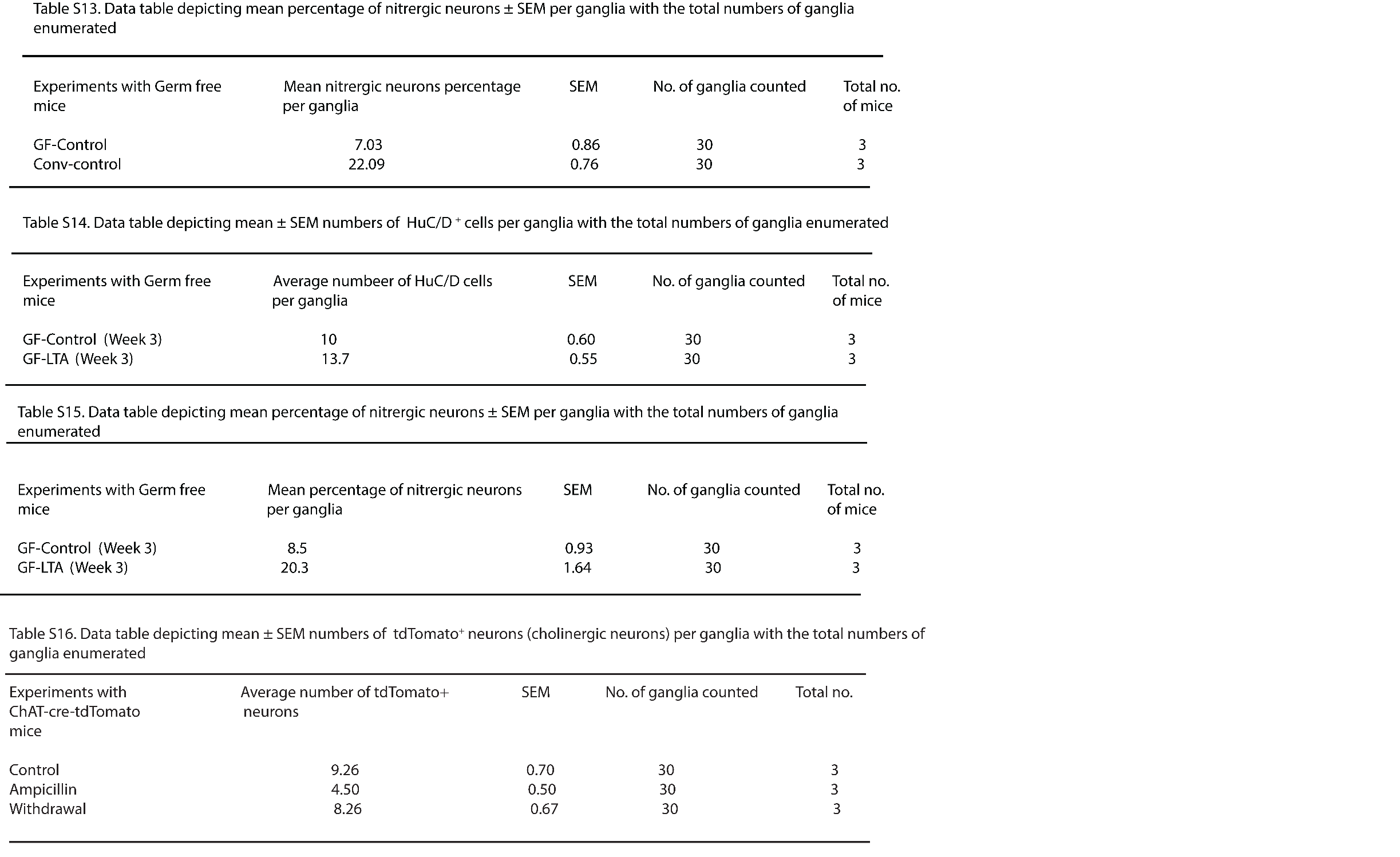

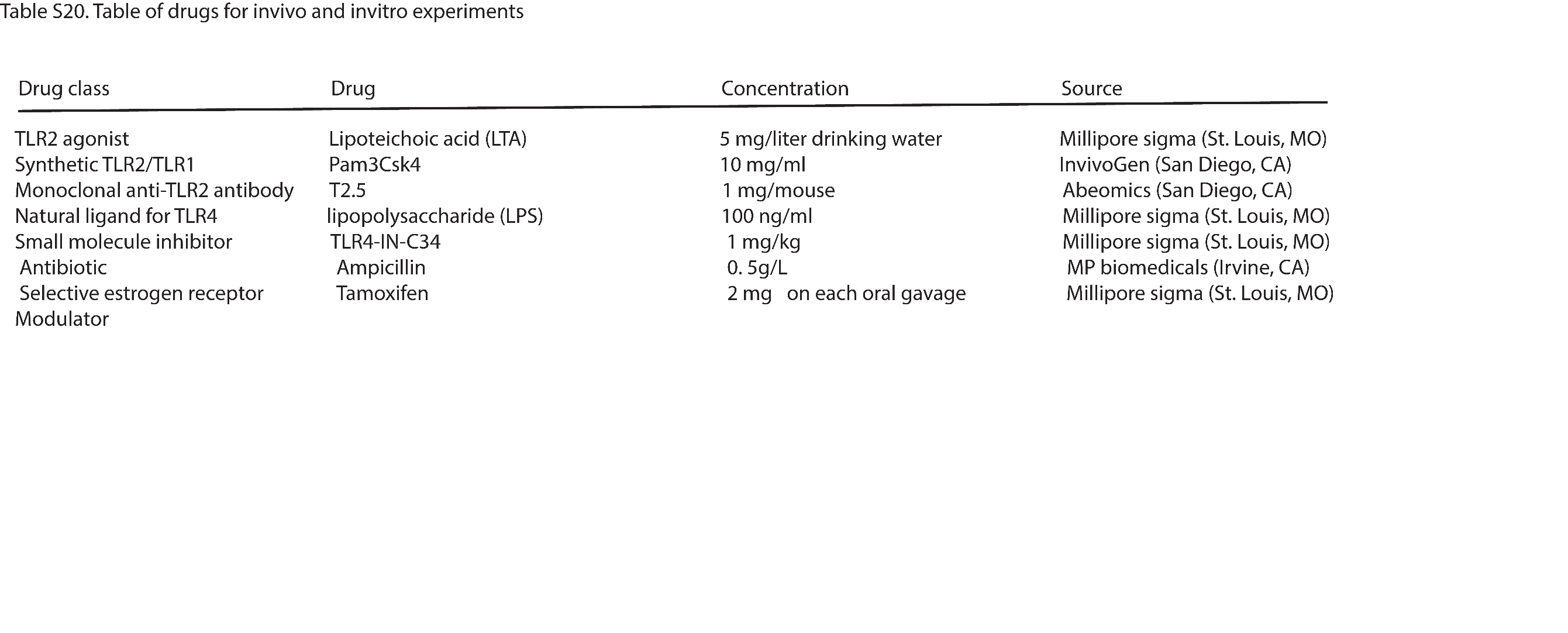

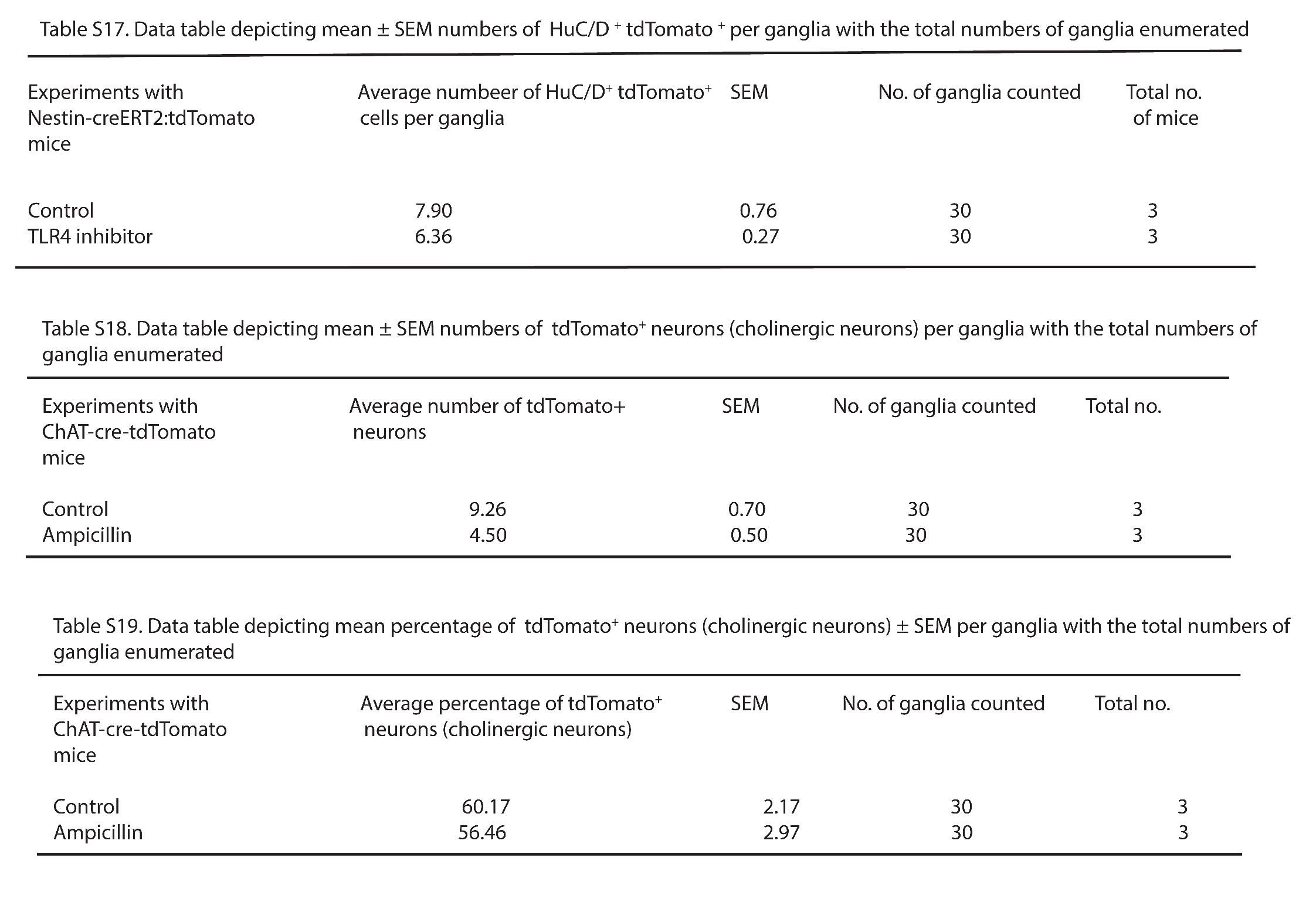

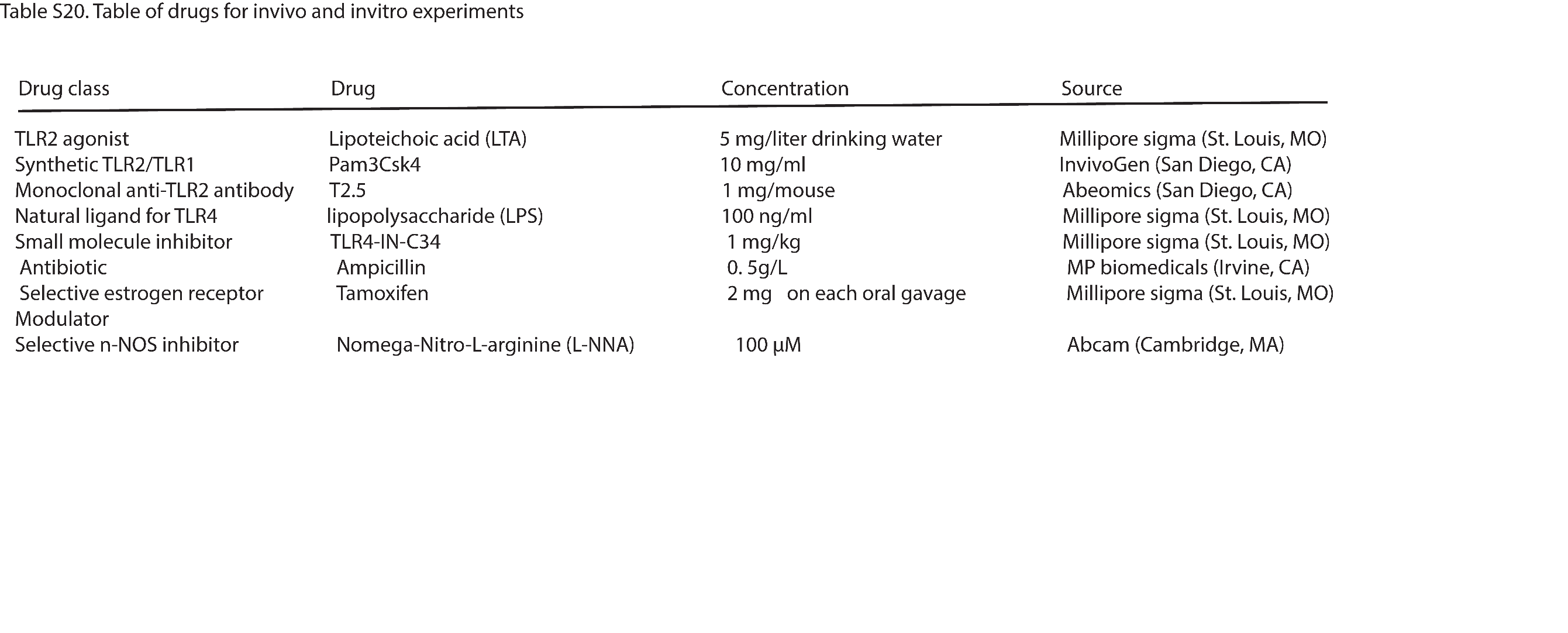

.

1.	Mignone JL, Kukekov V, Chiang AS, et al. Neural stem and progenitor cells in nestin-GFP transgenic mice. J Comp Neurol 2004;469:311-24.

2.	Anitha M, Vijay-Kumar M, Sitaraman SV, et al. Gut microbial products regulate murine gastrointestinal motility via Toll-like receptor 4 signaling. Gastroenterology 2012;143:1006-16 e4.

3.	Kozich JJ, Westcott SL, Baxter NT, et al. Development of a dual-index sequencing strategy and curation pipeline for analyzing amplicon sequence data on the MiSeq Illumina sequencing platform. Appl Environ Microbiol 2013;79:5112-20.

4.	Kulkarni S, Micci MA, Leser J, et al. Adult enteric nervous system in health is maintained by a dynamic balance between neuronal apoptosis and neurogenesis. Proc Natl Acad Sci U S A 2017.

5.	Becker L, Peterson J, Kulkarni S, et al. Ex vivo neurogenesis within enteric ganglia occurs in a PTEN dependent manner. PLoS One 2013;8:e59452.

6.	de Muinck EJ, Trosvik P. Individuality and convergence of the infant gut microbiota during the first year of life. Nat Commun 2018;9:2233.

7.	Ley RE, Hamady M, Lozupone C, et al. Evolution of mammals and their gut microbes. Science 2008;320:1647-51.

8.	Li K, Bihan M, Yooseph S, et al. Analyses of the microbial diversity across the human microbiome. PLoS One 2012;7:e32118.

9.	Bik EM, Eckburg PB, Gill SR, et al. Molecular analysis of the bacterial microbiota in the human stomach. Proc Natl Acad Sci U S A 2006;103:732-7.

10.	Callahan BJ, McMurdie PJ, Rosen MJ, et al. DADA2: High-resolution sample inference from Illumina amplicon data. Nat Methods 2016;13:581-3.

11.	Cole JR, Wang Q, Fish JA, et al. Ribosomal Database Project: data and tools for high throughput rRNA analysis. Nucleic Acids Res 2014;42:D633-42.

12.	Bodenhofer U, Bonatesta E, Horejs-Kainrath C, et al. msa: an R package for multiple sequence alignment. Bioinformatics 2015;31:3997-9.

13.	Schliep KP. phangorn: phylogenetic analysis in R. Bioinformatics 2011;27:592-3.

14.	Callahan BJ, Sankaran K, Fukuyama JA, et al. Bioconductor Workflow for Microbiome Data Analysis: from raw reads to community analyses. F1000Res 2016;5:1492.

15.	McMurdie PJ, Holmes S. phyloseq: an R package for reproducible interactive analysis and graphics of microbiome census data. PLoS One 2013;8:e61217.

16.	Fernandes AD, Reid JN, Macklaim JM, et al. Unifying the analysis of high-throughput sequencing datasets: characterizing RNA-seq, 16S rRNA gene sequencing and selective growth experiments by compositional data analysis. Microbiome 2014;2:15.
